# Rapid sensing and relaying of cellular hyperosmotic-stress signals via RAF–SnRK2 core condensates

**DOI:** 10.64898/2026.01.03.697504

**Authors:** Guting Liu, Zhen Lin, Guanquan Lin, Xinyong Wang, Xiaolei Liu, Zhaobo Lang, Jian-Kang Zhu, Pengcheng Wang

**Author notes:** These authors contribute equally to this work (Guting Liu, Zhen Lin). Correspondence (J.K.Z.), and (P.W.).

## Abstract

Hyperosmolarity caused by drought, high salinity, or cold stress inhibits plant growth and crop productivity^1,2^. A conserved protein-kinase cascade of cytosolic B-RAFs and SnRK2s is rapidly activated upon osmotic stresses to initiate downstream adaptive responses, which represents one of the fastest known responses to osmotic stress in plants^3–8^. How the kinase cascade is activated by osmotic stress is unknown. Here, we show that *Arabidopsis* B4 subgroup RAFs have intrinsically disordered regions and directly sense both ionic and non-ionic hyperosmolarity by reversible condensation. B4-RAFs recruit and co-condense with subclass-I SnRK2s to phosphorylate and turn on SnRK2s, evading the non-condensable inhibitory A-clade PP2C phosphatases. This straightforward osmosensing and relaying module can be fully reconstituted in *E. coli* by co-expressing three components or in solution in a test tube using recombinant proteins. Our findings identify B-RAFs as the chief cellular osmosensors that detect low water potential by co-condensation, forming a signal hub with SnRK2s to orchestrate adaptive responses in plants, and represent an evolutionarily conserved osmosensing mechanism across kingdoms.

## Introduction

Limited water availability is one of the most fundamental challenges that land plants must contend with. Drought, high salinity and low temperature cause hyperosmotic stress in cells. This suppresses plant growth — with major implications for crop yield.^1,2^ Upon exposure to hyperosmotic stress, B-subgroup RAPIDLY ACCELERATED FIBROSARCOMA (RAF) protein kinases are activated within seconds in plants,^3–8^ which represents one of the fastest known responses to osmotic stress in plants. B-RAFs then phosphorylate and activate plant-specific SNF1-RELATED PROTEIN-KINASE 2s (SnRK2s).^5,9–13^ Activated SnRK2s phosphorylate hundreds of downstream effectors, including transcription factors and transporters,^14,15^ to initiate adaptive stress responses such as stomatal closure, ^16,17^ growth inhibition,^18^ biosynthesis of compatible osmolytes, and the up-regulation of genes encoding protective proteins. ^19–21^

In the model plant *Arabidopsis thaliana*, 22 B-RAFs and 10 SnRK2s comprise a kinase cascade that orchestrates the signaling of osmotic stress and the ‘stress hormone’ abscisic acid (ABA).^9,11,22^ Within this cascade, B4-subgroup RAFs are activated earliest upon hyperosmolarity stimuli, and these in turn directly phosphorylate conserved serine/threonine residues in the activation loop to activate subclass-I SnRK2s (SnRK2.1/4/5/9/10).^4,5,13^ Hyperosmolarity-induced activation of these SnRK2s is completely abolished in *OK*^130^–*null* high-order *b4-raf* mutants, a mutant knocking-out all seven B4-subgroup *RAFs*, underscoring the central role of B4-RAFs in initiating the osmotic-stress signaling cascade.

By contrast, the B2 and B3 RAFs are required to activate ABA-dependent SnRK2.2/3/6 proteins upon ABA and hyperosmolarity.^9,11,23,24^ Under non-stress conditions, A-clade PP2C phosphatases dephosphorylate and inhibit SnRK2.2/3/6.^25–27^ ABA ligand binds to PYRABACTIN RESISTANCE 1 (PYR1)/PYR1-LIKE (PYL)/REGULATORY COMPONENTS OF ABA RECEPTOR (RCAR) receptors to inhibit PP2Cs, thereby releasing SnRK2s from inhibition.^28,29^ B-RAFs, which have basal activity, can then phosphorylate SnRK2s to allow ABA signaling to initiate.^11,30^ Though PP2Cs also dephosphorylate ABA-independent SnRK2s, activation of RAFs or these SnRK2s upon hyperosmotic stress occurs independently of ABA biosynthesis and receptor-coupled core signaling, which are unaffected in mutants of A-clade PP2Cs,^4,31^ ABA biosynthesis enzymes,^32^ or ABA receptors.^33^ Activation of this RAF–SnRK2 cascade is also independent of the membrane osmosensor OSCA1^4,34^. How this core kinase cascade is initially turned on by osmotic stress remains unknown.

## RESULTS

### B4-RAFs sense hyperosmolarity by condensation

Our identification of conserved intrinsically disordered regions (IDRs)^8^ in B-RAFs (Figure S1A ) hint at the potential involvement of condensates in B-RAF activation. Supporting this notion, application of 800 mM mannitol, a typical hyperosmolarity treatment used by related studies,^4,5,32,35,36^ to Arabidopsis plants rapidly induces condensation of native-promoter-driven RAF40–GFP or RAF24–GFP in the *OK^1^*^30^*-null* background, and GFP–RAF20 in the wild-type Col-0 background, within 1 min *in vivo* (Figures 1A and 1B). RAF8–GFP and GFP–RAF11 also form osmotic-stress-induced punctate structures (Figure S1B). No condensation of free GFP in *35S_pro_:GFP* transgenic plants or recombinant GFP/YFP/RFP alone is observed under the same conditions (Figures S1C, S1D). Neither cold nor heat treatment induced condensation of RAF40–GFP (Figure S1E). Intriguingly, mild hyperosmolarity, i.e. 300 or 500 mM mannitol treatment, induced the condensation of RAF24–GFP, but not GFP–RAF20 and RAF40–GFP within 5 min in transgenic plants (Figures 1C, S2A, and S2B ), suggesting different B4-RAF members may vary in sensitivity to hyperosmolarity conditions. Pre-incubation with 1,6-hexanediol (1,6-HD, a chemical that disrupts global condensates)^37^ abolishes mannitol-induced RAF40–GFP and RAF24–GFP condensation *in planta* (Figure 1D). The mannitol-triggered condensation of RAF40–GFP and could also be reversed by transferring *RAF40*–*GFP* seedlings back into half-strength Murashige and Skoog liquid control medium (referred to as Mock) (Figure 1E ). Mannitol-triggered RAF40–GFP and RAF24–GFP condensates are reformed by a second hyperosmotic stress treatment (Figure 1E). Hyperosmotic stress triggers a mobility shift for RAF24–GFP, serving as a readout of its phosphorylation and activation.^38^ Consistent with reversible condensation, the RAF24–GFP mobility shift could also be abolished by withdrawing mannitol and reinduced by a second round of treatment (Figure S2C, top panel). Like RAF24–GFP, the mobility shift of RAF35 was also abolished by withdrawing mannitol and re-induced by a second round of treatment, as indicated by immunoblot with an antibody recognizing native RAF35.^38^ We observed the phosphorylation of a conserved phosphosite corresponding to Ser166 in SnRK2.4 and Ser175 in SnRK2.6,^24^ which is the target site of B-RAFs.^4,9,11^ Like the RAF24–GFP and RAF35–GFP mobility shift, this phosphorylation of SnRK2s is consistent with the condensation of RAF24–GFP (Figures 1F, S2C, and S2D), suggesting the condensation of B4-RAFs correlates with their activation upon hyperosmolarity.

**Figure 1.**
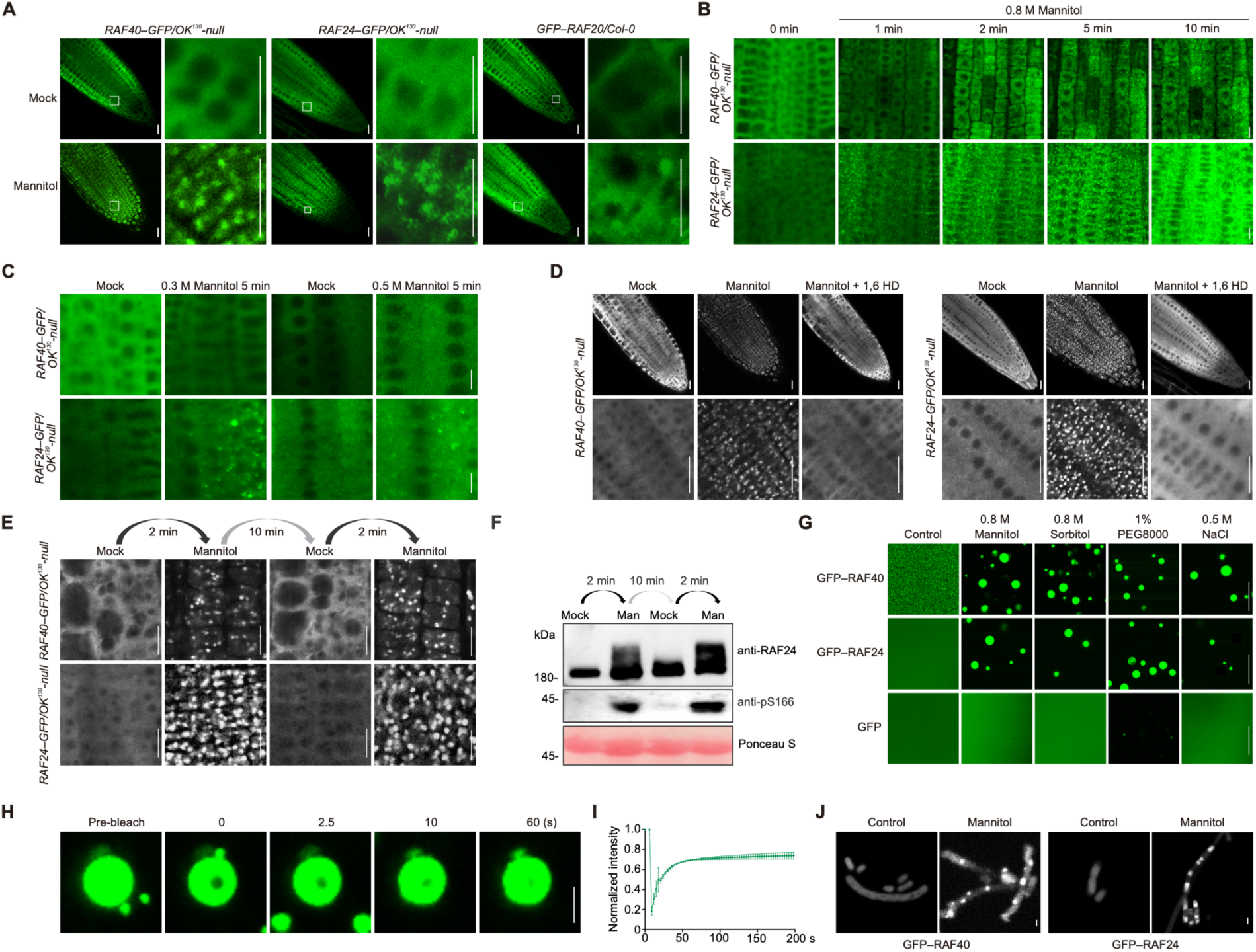
B4 RAFs undergo condensation upon hyperosmotic stresses. (**A**) Mannitol treatment triggers condensation of RAF40*–*GFP (left), RAF24*–*GFP (middle), and GFP*–*RAF20 (right) in transgenic Arabidopsis plants with the indicated genetic backgrounds. The small white boxes in the left-hand images are zoomed in at right. (**B**) Distribution dynamics of RAFs in root-tip cells of *RAF40_pro_:RAF40–GFP/OK^130^-null* and *RAF24_pro_:RAF24–GFP/OK^130^-null* transgenic plants upon hyperosmotic (1/2 MS + 800 mM mannitol) treatment. (**C**) Differential responses of RAF40*–*GFP and RAF24*–*GFP under mock (1/2 MS) or hyperosmotic stress (1/2 MS + 300 or 500 mM mannitol) treatments. (**D**) Mannitol-trigged RAF40*–*GFP and RAF24*–*GFP condensates are abolished by the application of 1,6-HD. (**E**) The mannitol-triggered condensation of RAF40*–*GFP and RAF24*–*GFP is reversible. (**F**) The mannitol-induced changes in RAF24 mobility are reversible. Immunoblotting shows the mobility shift of RAF24 (top), phosphorylation of Ser166 (a conserved B-RAF target site corresponding to Ser166 in SnRK2.4, middle), and loading control of Ponceau S staining (bottom). (**G**) Various hyperosmotic treatments trigger condensation of recombinant GFP*–*RAF40 and GFP*–*RAF24, but not free GFP proteins, *in vitro*. (**H**) FRAP of GFP*–*RAF40 condensates formed *in vitro*. Time 0 s indicates the time of the photobleaching pulse. (**I**) Time course of recovery after photobleaching GFP*–*RAF40 condensates. (**J**) Representative confocal-microscopy images of *E. coli* cells expressing GFP*–*RAF40 and GFP*–*RAF24. Cells were treated without or with 800 mM mannitol for 5 min. Scale bars: 5 μm in (**A–E, G**); 1 μm in (**H**) and (**J**). See also Figure S1–S3.

We next examined the ability of recombinant B4-RAFs to form punctate structures *in vitro*. We could only obtain recombinant full-length GFP–RAF40, but not RAF40–GFP or any other tagged B4-subgroup RAF, in *E. coli* with standard lysogeny broth (LB) medium with 10 g/L (about 171 mM) NaCl. The macromolecular-crowding reagent PEG8000 at 10% is sufficient to induce condensation of 0.1 µM GFP–RAF40, while 0.1% PEG8000 is sufficient to trigger condensation of 5 µM GFP–RAF40 (Figure S3). Higher concentrations of PEG8000, e.g., 10%, caused the segregation of recombinant GFP–RAF40 within 1 min (Figure S3A). Interestingly, treatment with relatively high concentration of salt, like LiCl, KCl, or NaCl, or low-molecular-weight chemicals, like mannitol, ethylene glycol (EG), and PEG200, also induces condensation of recombinant GFP–RAF40 in solution (Figures 1G, S3B, and S3C). Fluorescence recovery after photobleaching (FRAP) assays showed that GFP–RAF40 undergoes liquid-like droplets in solution (Figures 1H and 1I, Video S1).

Using LB with 2.5 g/L (about 43 mM) NaCl, which may prevent the hyperosmolarity-induced segregation of recombinant RAFs during cell culture (Figure S3D), we could obtain soluble recombinant GFP–RAF24 and GFP–RAF20 (Figure S3D, Lanes 3 and 7). Recombinant GFP–RAF24 and GFP–RAF20 also condense in solution with mannitol, sorbitol, PEG8000, and NaCl (Figures 1G, S3E, and S3F). Because 800 mM mannitol is commonly used for rapid RAF and SnRK2 activation *in planta* ^4,5,32,35,36^ and could trigger condensation of recombinant B4-RAFs here and for the IDR of SEUSS ^39^ (Figure S3G), we chose to use 800 mM mannitol and 5 µM recombinant proteins for most assays hereinafter. Effects with 800 mM mannitol are reproducible with 1% PEG8000 in all tested experiments. Such hyperosmotic-stress-induced condensates appear even in *E. coli* after adding 800 mM mannitol to the medium (Figure 1J).

These data suggest that the sequence and/or property(s) of B4-RAFs, represented by RAF40/RAF24/RAF20 here, define the ability to form condensates in solution upon both ionic and non-ionic hyperosmotic stress.

Fluorescence of RAF40 condensates does not overlap with the processing-body (p-body) marker DCP1,^40^ stress-granule maker RBP47b–mCherry,^40^ or the osmosensor DCP5 ^41^ (Figures S4A–C), in both transgenic Arabidopsis or tobacco leaves (Figure S4D). Unlike RAF18, which localizes in p-bodies,^5^ both RAF40–GFP and SnRK2.4–BFP do not appear to colocalize with DCP1–mCherry under these conditions (Figures S4D and 4E).

### Multiple domains are required for B4-RAF condensation

B4-subgroup RAFs contain a PB1 (Phox and Bem1) domain at the N-terminus that confers protein–protein interactions (designated ‘NT’, residues 1–272 in RAF40), three IDRs (‘IDR’, residues 273–812 in RAF40), and a C-terminal kinase domain (‘KD’, residues 813–1,117 in RAF40) (Figure 2A). To determine which RAF40 region may be required for hyperosmolarity-induced condensation, we generated truncations carrying the specified regions fused to GFP (Figure 2A). Without mannitol, GFP–RAF40^IDR^, but not the full-length or any other truncated version, has slight punctate structures in solution (Figure 2B). Full-length GFP–RAF40 and variants containing IDRs condense in a mannitol-dependent manner. Variants with GFP–RAF40^IDR^, but not GFP–RAF40^NT^ or GFP–RAF40^KD^, are sufficient to drive their condensation upon mannitol treatment (Figure 2B). However, the NT and KD may contribute to the condensation of GFP–RAF40, as the GFP–RAF40^IDR–KD^ and GFP–RAF40^NT–IDR^ condensates differ from full-length GFP–RAF40, in the context of size/intensity of condensates under the same conditions (Figures 2B and 2C). Similarly, we observed that the IDR is essential for, and the NT and KD contribute to, the condensation of RAF40–GFP in transgenic Arabidopsis seedlings expressing different RAF truncations in the *OK^1^*^30^*-null* mutant background (Figure 2D).

**Figure 2.**
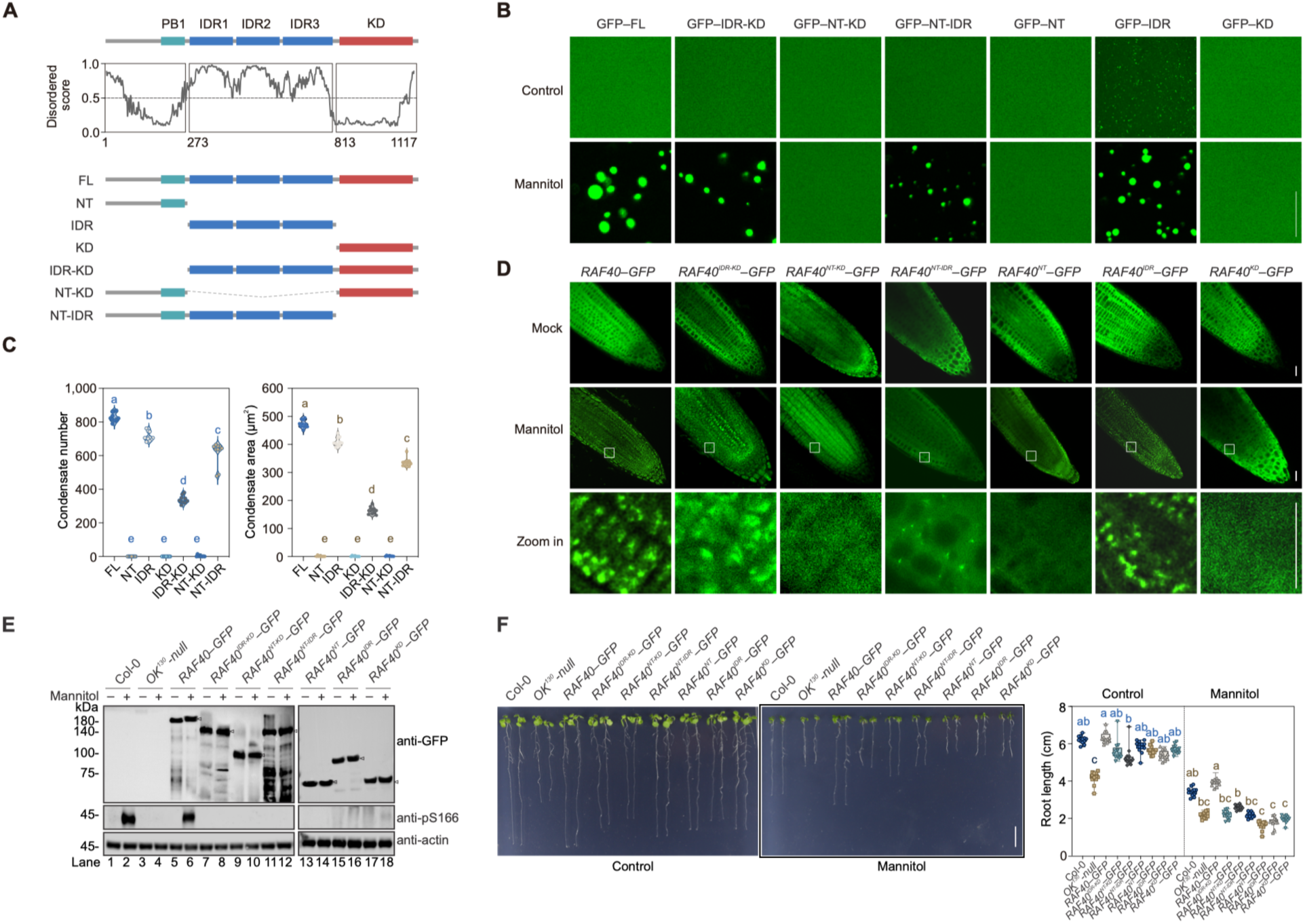
The IDR of RAF40 is responsible for its condensation and is required for osmotic-stress tolerance. (**A**) Top, RAF40 domain organization. Middle, RAF40 intrinsically disordered regions predicted using IUPred2. Bottom, diagram of truncated RAF40 variants. (**B**) *In vitro* condensation assay of 5 µM recombinant GFP-fused RAF40 variants. (**C**) Quantification of condensate number and area of GFP-fused RAF40 variants in buffer containing 800 mM mannitol as in **(B)** (*p* < 0.0001, one-way ANOVA followed by Tukey’s post-hoc test). n = 9 independent replicates. (**D**) Condensation assay in transgenic Arabidopsis complementation lines carrying GFP-fused RAF40 variants in the *OK^130^-null* background. (**E**) Immunoblots of RAF40*–*GFP abundance (top) and phosphorylation of a conserved serine corresponding to Ser166 in SnRK2.4 (middle) in different *OK^130^-null* complementation lines characterized in (**D**). Arrows on the anti-GFP blot indicate the GFP fused RAF40 variants. Actin was used as a loading control. (**F**) Photographs (left) of 7-d-old seedlings of Col-0, *OK^130^-null,* and different complementation lines grown on 1/2 MS medium (Mock) or 1/2 MS medium with 200 mM mannitol. Right panel, primary root length of seedlings shown at left. (*p* < 0.05, one-way ANOVA followed by Tukey’s post-hoc test). For (**B**), representative images of n = 9 independent experiments; (**D**), (**E**), and (**F**), n = 3 independent experiments. Scale bars: 5 μm in (**B** and **D**); and 1 cm in (**F**). See also Figure S4.

We then asked whether B4-RAF condensation is important for their activation upon hyperosmotic stress. Mannitol treatment quickly induces the activation of RAF40–GFP in the Col-0 background and *RAF40*–*GFP*/*OK^1^*^30^*-null* plants, but not in the negative control *OK^1^*^30^*-null* mutant, as indicated by immunoblotting with anti-pS166 antibodies (Figure 2E, lanes 1–6, middle panel). Deletion of IDR, NT or KD strongly abolishes or impairs mannitol-triggered Ser166 phosphorylation in the complementation lines expressing GFP fusions (Figure 2E, lanes 7–18, middle panel). Full-length RAF40–GFP, but not RAF40^IDR–KD^, RAF40^NT–KD^, RAF40^NT–IDR^, RAF40^NT^, RAF40^IDR^, or RAF40^KD^ variants, could complement the hypersensitive growth-inhibition phenotype of *OK^1^*^30^*-null* seedlings to mannitol and NaCl treatments (Figures 2F and S4F).

### Q-to-A mutation abolishes B4-RAF condensation

We further attempted to abolish or impair RAF40 condensation by mutation of ‘sticker’ or ‘spacer’ amino acids within IDRs, without deletion of large fragments (Figure S5A). RAF40 has 36 glutamine (Q), 27 arginine (R), and 15 tyrosine (Y) residues in the IDR of RAF40 and these are considered spacer or sticker amino acids^42,43^. The recombinant protein carrying 36 glutamine (Q) to alanine (A) mutations, RAF40^QA^, has abolished mannitol-triggered condensation *in vitro* (Figures 3A and 3B). However, variants carrying 15 tyrosine to phenylalanine mutations, RAF40^YF^, or carrying 27 arginine to lysine mutations, RAF40^RK^, keep their ability to condense in solution with mannitol (Figure S5B). RAF40^QA^–GFP lost the ability to form condensates in *RAF40^QA^*–*GFP*/*OK^1^*^30^*-null* transgenic plants (Figures 3C and 3D). The RAF40^QA^–GFP construct only partially complements the mannitol- and NaCl-hypersensitivity phenotypes of *OK^1^*^30^*-null* in the context of root growth (Figures 3E and 3F). Consistently, the mannitol-triggered phosphorylation of the conserved serine residue, corresponding to Ser166 in SnRK2.4, is also impaired in *RAF40^QA^*–*GFP*/*OK^1^*^30^*-null*, compared to *RAF40*–*GFP*/*OK^1^*^30^*-null* (Figure 3G, middle panel), though the abundance of RAF40–GFP in these lines is comparable (Figure 3G, top panel). Thus, both the kinase activity and condensation of RAF40 are required for its activation by osmotic stress. It is notable that the activation of B4-RAFs and SnRK2s are independent of OSCA1^21,34^ and DCP5^41^ (Figure S5C).

**Figure 3.**
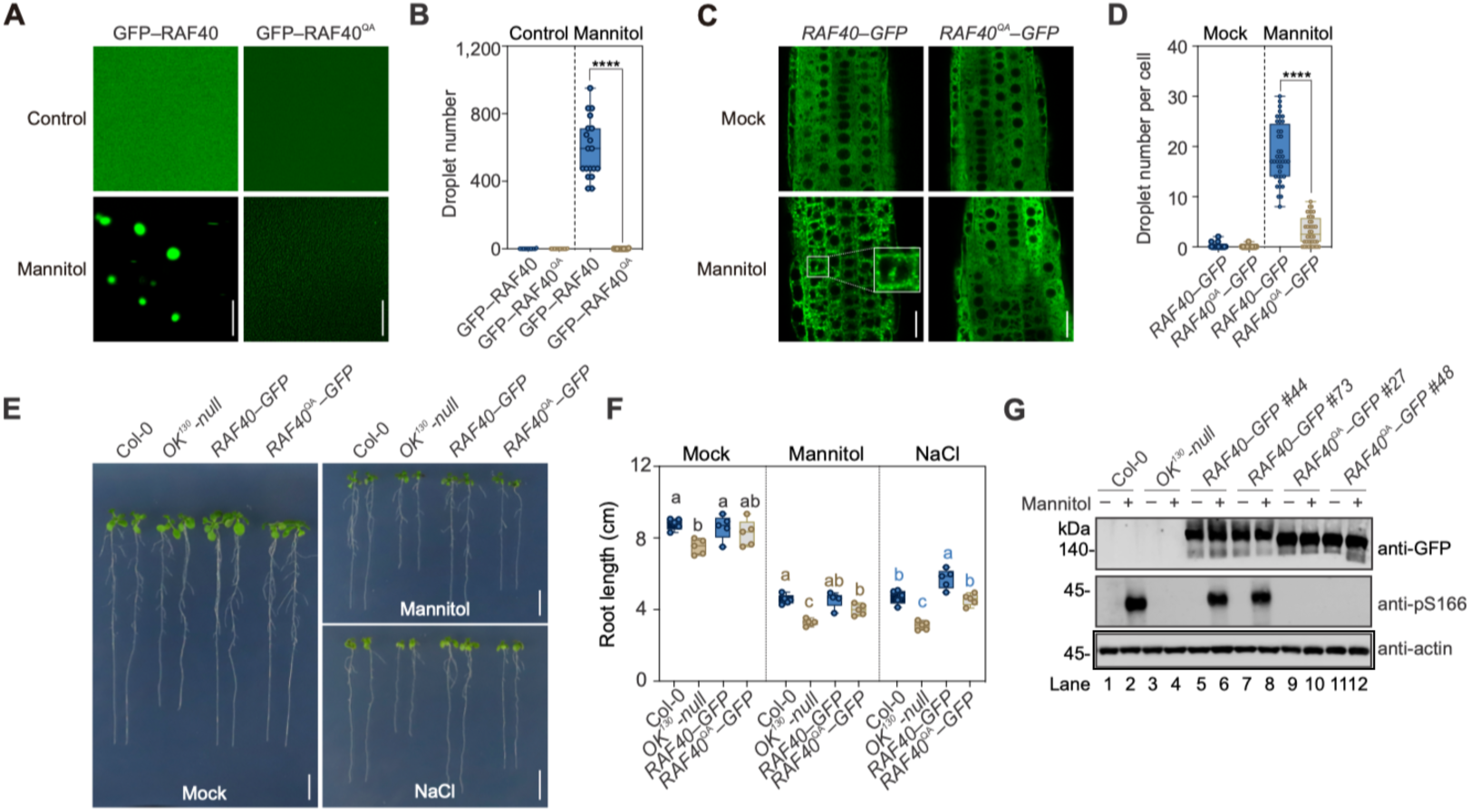
Mutation of 36 glutamine residues in RAF40 abolishes osmotic-stress-induced condensation and alters stress responses. (**A**) Hyperosmotic treatment triggers condensation of recombinant GFP*–*RAF40, but not GFP*–*RAF40^QA^ proteins *in vitro*. (**B**) Quantification of condensate number of recombinant GFP*–*RAF40 and GFP*–*RAF40^QA^ proteins in buffer without or with 800 mM mannitol as in **(A)**. ****, *p* < 0.0001, unpaired two-tailed *t*-test. Data are mean ± SD of n = 9 independent replicates. (**C**) Condensation assay in *RAF40–GFP/OK^130^-null* and *RAF40^QA^–GFP/OK^130^-null* complementation lines. (**D**) Quantification of RAF40 condensates per cell in **(C)**. ****, *p* < 0.0001, unpaired two-tailed *t*-test. Data are mean ± SD of n = 9 independent replicates. (**E**) Photographs of 10-day-old seedlings of Col-0, *OK^130^-null, RAF40–GFP/OK^130^-null,* and *RAF40^QA^–GFP/OK^130^-null* complementation lines grown on 1/2 MS or with 200 mM mannitol or 100 mM NaCl. (**F**) Primary root length of seedlings shown in **(E)**. Different letters (a–c) indicate statistically significant differences between groups (*p* < 0.05, one-way ANOVA followed by Tukey’s post-hoc test). (**G**) Immunoblots of RAF40*–*GFP and RAF40^QA^*–*GFP abundance (top) and phosphorylation of a conserved serine corresponding to Ser166 in SnRK2.4 (middle) in (**E**). Actin was used as a loading control. Scale bars: 5 μm in (**A**) and (**C**); 1 cm in (**E**). (**C**), (**E**), and (**G**), n = 3 independent experiments. See also Figure S5.

### B4-RAFs recruit subclass-I SnRK2 to form co-condensates from PP2Cs

Next, we assessed whether SnRK2s themselves, as B-RAF targets, may have IDRs and undergo condensation upon hyperosmotic stress treatment. The ABA-independent or subclass-I SnRK2s (namely SnRK2.1, SnRK2.4, SnRK2.5, SnRK2.9 and SnRK2.10)^44^ have short IDRs predicted in their C termini (Figures 4A, 4B, and S6A), but the ABA-dependent subclass-III members SnRK2.2, SnRK2.3, and SnRK2.6 lack them (Figures 4A, 4B, and S6A). ABA-inducible subclass-II SnRK2.7 and SnRK2.8 have ambiguous disorder scores (Figures 4A and S6A). Consistent with a recent study^45^, recombinant SnRK2.4–YFP, SnRK2.1–YFP, SnRK2.5–YFP, and SnRK2.9–YFP show condensation upon 800 mM mannitol and 5 µM recombinant proteins (Figures 4C, left panel, and S6B). SnRK2.10–YFP only forms condensates with 800 mM mannitol and 50 µM recombinant proteins (Figure S6C). Deletion of its C-terminal IDR (303–371 aa), but not its KD (1–260 aa), abolishes this condensation of recombinant SnRK2.4–YFP (Figure 4C, central and right panels). In transgenic plants expressing native-promoter-driven *SnRK2.4–GFP*, mannitol treatment also triggers SnRK2.4–GFP condensation (Figure 4D). In contrast, SnRK2.6 purified from *E. coli* did not condense in solution with 800 mM mannitol (Figure S6D), nor was condensation observed in *SnRK2.6*–*GFP* plants^18^ (Figure S6E).

**Figure 4.**
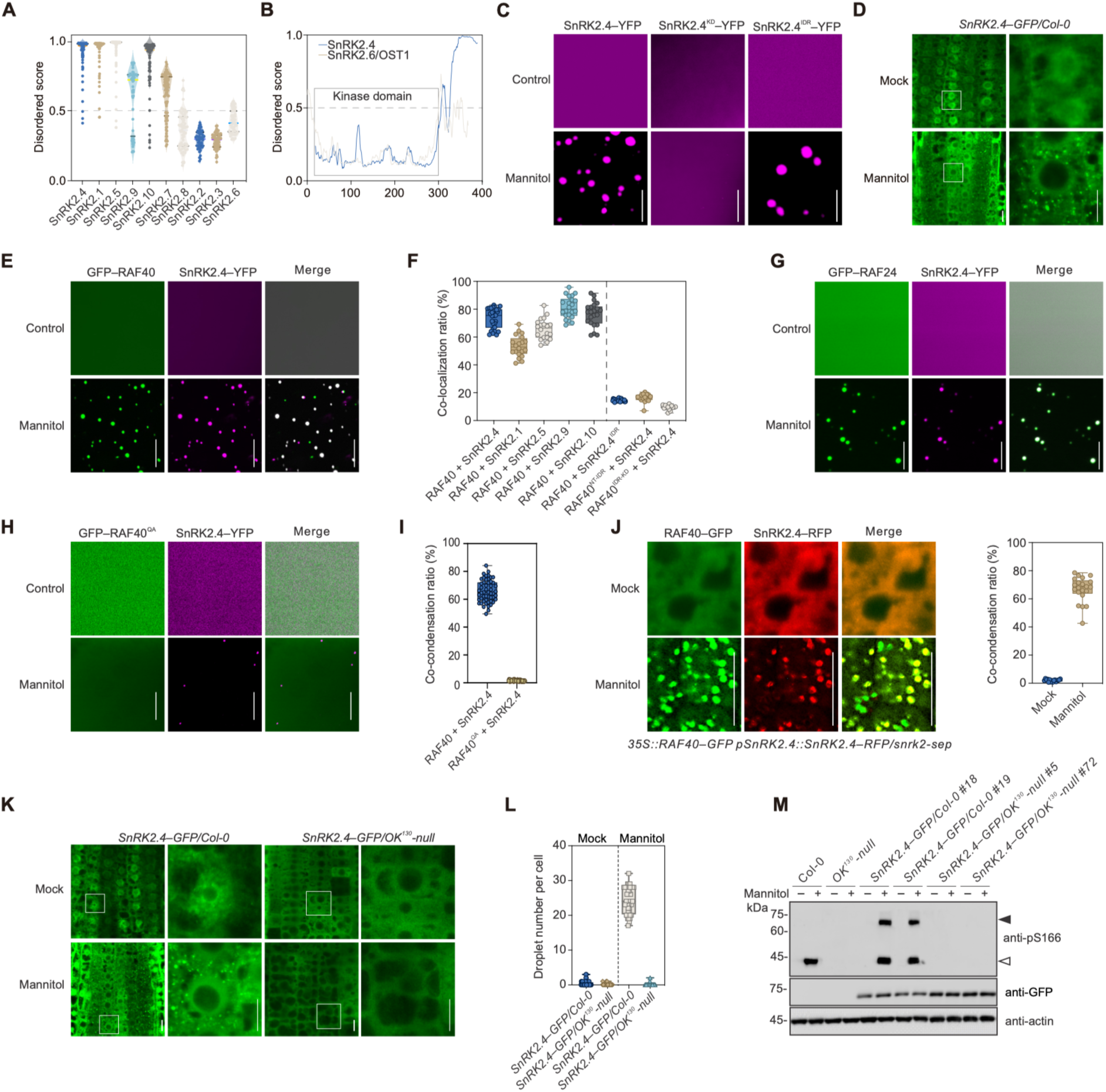
RAF40 and RAF24 co-condense with SnRK2s under hyperosmolarity. (**A**) The intrinsically disordered score of SnRK2 C-terminal regions predicted by IUPred2. (**B**) The intrinsically disordered regions of full-length SnRK2.4 and SnRK2.6 predicted by IUPred2. (**C**) *In vitro* condensation assay with 5 µM recombinant full-length and truncated SnRK2.4–YFP proteins. (**D**) Condensation assay of SnRK2.4–GFP/Col-0 under 1/2 MS (Mock) or with 800 mM mannitol. Images at right are zoomed in from the white boxes at left. (**E**) *In vitro* condensation assay showing the co-condensation GFP–RAF40 and SnRK2.4–YFP in solution without or with 800 mM mannitol. (**F**) Quantification of colocalization ratio between GFP-fused RAF40 variants and YFP-fused SnRK2.4 variants in buffer containing 800 mM mannitol as in **(E)**. (**G**) *In vitro* condensation assay showing the co-condensation GFP–RAF24 and SnRK2.4–YFP in solution without or with 800 mM mannitol. (**H**) *In vitro* condensation assay of GFP–RAF40^QA^ and SnRK2.4–YFP in solution without or with 800 mM mannitol. (**I**) Quantification of colocalization ratio between SnRK2.4–YFP and GFP–RAF40, or SnRK2.4–YFP and GFP–RAF40^QA^ in control buffer or with 0.8 M mannitol treatment. (**J**) Colocalization of fluorescence of RAF40–GFP and SnRK2.4–RFP in *35S_pro_:RAF40*–*GFP SnRK2.4_pro_:SnRK2.4*–*RFP/snrk2-sep* transgenic plants. Quantification of colocalized puncta formed upon mannitol treatment is shown at right. (**K**) Fluorescence of SnRK2.4–GFP in *SnRK2.4–GFP/*Col-0 and *SnRK2.4*–*GFP/OK^130^-null* plants in 1/2 MS (Mock) or with 800 mM mannitol. Images at right are zoomed in from the white boxes at left. (**L**) Quantification of puncta per cell in *SnRK2.4*–*GFP*/Col-0 and *SnRK2.4*–*GFP/OK^130^-null* transgenic lines, as shown in **(K)**. (**M**) Immunoblots of SnRK2.4–GFP abundance (middle) and phosphorylation of conserved serine residue corresponding to Ser166 in SnRK2.4 (top) in *SnRK2.4*–*GFP*/Col-0 and *SnRK2.4*–*GFP/OK^130^-null* transgenic lines. The open and closed arrows indicate the endogenous SnRK2s and transgenic SnRK2.4–GFP, respectively. Actin was used as a loading control. For (**C**), (**E**), (**G**), and (**H**), representative images of n = 7 independent experiments; (**D**), (**J**), (**K**), and (**M**), representative images of n = 3 independent experiments. Scale bars: 5 μm in (**C**), (**E**), (**G**), and (**H**); and 10 μm in (**D**), (**J**), and (**K**). See also Figure S6 and S7.

We then tested if RAF40 and SnRK2.4 can co-condense together *in vivo* and *in vitro*. Recombinant GFP–RAF40 and SnRK2.4–YFP were mixed and upon exposure to 800 mM mannitol, their fluorescence largely overlapped within condensates (Figures 4E and 4F). Additionally, GFP–RAF40 also co-condenses with SnRK2.1–YFP, SnRK2.5–YFP, SnRK2.9–YFP, SnRK2.10–YFP (Figures 4F and S7A). GFP–RAF24 also largely co-condenses with recombinant SnRK2.4–YFP in the presence of 800 mM mannitol (Figure 4G). Interestingly, the SnRK2.4 KD is essential for its co-condensation with RAF40–GFP. Almost no overlap was observed between GFP–RAF40 and SnRK2.4^IDR^–YFP condensates (Figures 4F and S7B). Deletion of the KD or NT region of RAF40 (GFP–RAF40^NT–IDR^ or GFP–RAF40^IDR–KD^) also abolishes its co-condensation with SnRK2.4–YFP (Figures 4F, S7C, and S7D). While the IDRs are sufficient for the condensation of RAF40 and SnRK2.4, the other regions of these proteins are necessary for their co-condensation. GFP–RAF40^QA^ did not detectably form condensates upon mannitol treatment, even in the presence of SnRK2.4–YFP (Figures 4H and 4I), and also abolishes its co-condensation with SnRK2.4–YFP (Figures 4H and 4I). The addition of GFP–RAF40^QA^ strongly impairs the condensation of SnRK2.4–YFP (Figures 4H and S7E).

We also examined the co-condensation of RAF40 and SnRK2.4 in plants. RAF40–GFP and SnRK2.4–RFP show about 70% of fluorescence overlap in the *35S_pro_*:*RAF40*–*GFP SnRK2.4*–*RFP/snrk2-sep* background, a transgenic line carrying native-promoter-driven *SnRK2.4* and over-expressing *RAF40–GFP*, with knock-out of native *SnRK2.1*, *SnRK2.4*, *SnRK2.5*, *SnRK2.7*, *SnRK2.8*, *SnRK2.9* and *SnRK2.10* (Figure 4J). Some non-overlapping punctate structures are likely because other native RAFs or SnRK2s compete with the tagged proteins to participate in the condensates upon hyperosmolarity. SnRK2.4–GFP did not form detectable condensates in *OK^1^*^30^*-null* seedlings that lack all seven *B4-RAFs* (Figures 4K and 4L), even though the *SnRK2.4*–*GFP/OK^1^*^30^*-null* lines have similar SnRK2.4–GFP abundance to that in *SnRK2.4*–*GFP*/Col-0 (Figure 4M). Consistent with the condensation result, Ser166 phosphorylation of both native SnRK2s and SnRK2.4–GFP is strongly induced by mannitol treatment in Col-0 and *SnRK2.4*–*GFP*/Col-0 seedlings, but not in *OK^1^*^30^*-null* and *SnRK2.4*–*GFP/OK^1^*^30^*-null* seedlings (Figure 4M, upper panel). Thus, RAF40 co-condenses with SnRK2.4 to form core condensates upon hyperosmotic stress treatment, leading to SnRK2.4 activation, and the condensation of SnRK2.4 *in vivo* is dependent on B4-RAF. RAF40–GFP retains its ability to condense under mannitol treatment in *RAF40–GFP/snrk2-sep* plants (Figure S7F), suggesting the condensation of RAF40–GFP is independent of presence of subclass-I SnRK2s.

### B4-RAFs recruit subclass-I SnRK2s

To phosphorylate and activate SnRK2s, RAF kinases must extricate SnRK2s from PP2C-mediated inhibition. To explore how RAF condensation affects this inhibition, we measured the condensation of A-clade PP2Cs *in vitro* and *in vivo*. Of the nine members of the A-clade, three of them (AHG1, ABI2, and HAI2) have IDRs in their N-termini with moderate scores. The remaining six have no obvious IDR (Figure S8A). We examined three transgenic lines carrying PP2Cs fused to fluorescent proteins: *35S_pro_:ABI2*–*GFP/Col-0*,^46^ *HAB1_pro_:HAB1–Venus*/Col-0,^47^ and *ABI1_pro_:ABI1–GFP*/*abi1 abi2*.^48^ None of them exhibit detectable condensation upon 800 mM mannitol treatment (Figure S8B), and assays in mesophyll protoplasts showed that all nine transiently expressed A-clade PP2Cs do not condense upon mannitol treatment (Figure S8C). We chose HAB1 as a case study to mimic signal processing in plants. SnRK2.4–YFP and HAB1–RFP were mixed ± RAF40–GFP, and condensation of these proteins is observed before and after mannitol treatment. As PP2C can inhibit at least 100-fold excess SnRK2s,^18,27^ we used 0.5 μM recombinant HAB1–RFP and 5 μM recombinant SnRK2.4–YFP, a ratio sufficient for HAB1 to abolish SnRK2.4 activity (Figures S9A and S9B ). HAB1–RFP impairs the mannitol-triggered condensation of SnRK2.4–YFP as reflected by the number and size of SnRK2.4–GFP condensates (Figure 5A). Addition of GFP–RAF40, but not GFP–RAF40^QA^, dramatically rescues the condensation of SnRK2.4–YFP in the presence of HAB1 (Figures 5B–5E). When all three proteins are present, mannitol treatment rapidly promotes the formation of SnRK2.4–YFP and GFP–RAF40 co-condensates, whereas HAB1–RFP remains soluble (Figures 5B and 5C). We obtained similar results with the substitution of GFP–RAF40 with GFP–RAF20 and GFP–RAF24 (Figures S9C–E), or the substitution of HAB1–RFP with the other two tested PP2Cs, HAB2–RFP and PP2CA–RFP (Figures S9F and S9G). Thus, hyperosmolarity-induced core condensates form for B-RAFs and SnRK2s, which allows SnRK2s to be spatially separated from A-clade PP2Cs, and evade inhibition by these PP2Cs. This may explain why osmotic-stress-triggered SnRK2 activation is independent of PP2Cs or other components in ABA-receptor-coupled core signaling.

**Figure 5.**
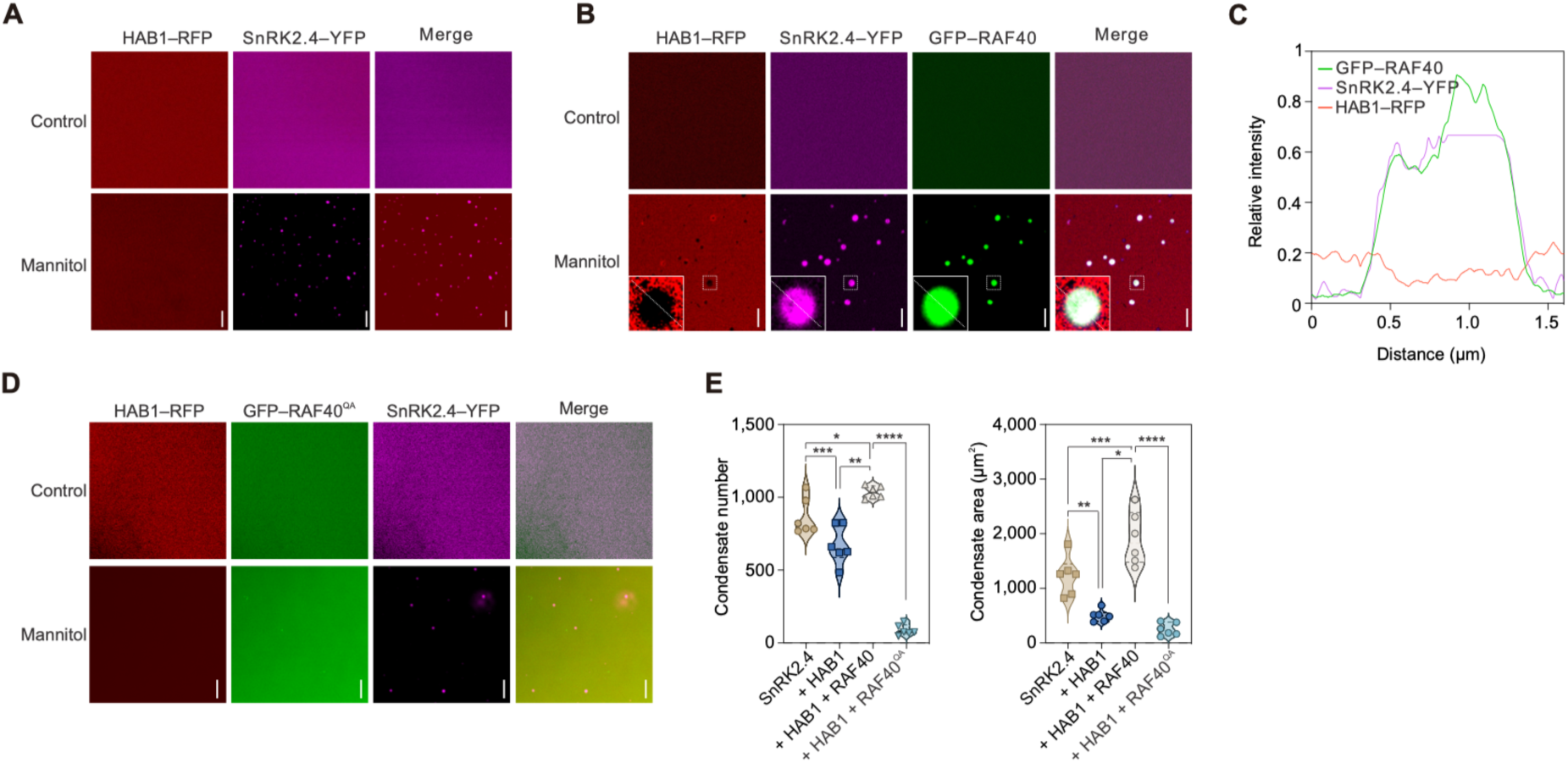
RAF40 recruits SnRK2.4 from HAB1 under hyperosmolarity. (**A**) *In vitro* condensation assay of 5 µM recombinant SnRK2.4–YFP with 0.5 µM HAB1–RFP. (**B**) *In vitro* condensation assay of 5 µM recombinant SnRK2.4–YFP with 0.5 µM HAB1–RFP and 5 µM GFP–RAF40. Insets for the mannitol treatment are zooms of the indicated small white boxes. (**C**) Relative intensity of SnRK2.4–YFP, HAB1–RFP, and GFP–RAF40 fluorescence from condensates marked in (**B**). (**D**) *In vitro* condensation assay of 5 µM recombinant SnRK2.4–YFP with 0.5 µM HAB1–RFP and 5 µM GFP–RAF40^QA^. (**E**) Quantification of the number and size of SnRK2.4–YFP puncta formed by SnRK2.4–YFP alone, SnRK2.4–YFP incubated with HAB1–RFP, SnRK2.4–YFP incubated with HAB1–RFP and GFP–RAF40, and SnRK2.4–YFP incubated with HAB1–RFP and GFP–RAF40^QA^ under mannitol treatment. ****, *p* < 0.0001, unpaired two-tailed *t-*test between two groups. Data are mean ± SD of n = 9 independent replicates. Scale bars: 5 μm in (**A**), (**B**) and (**D**). See also Figure S8 and S9.

### Reconstitution of RAF–SnRK2 osmosensing core module

We attempted to reconstitute this plant osmosensing core module in *E. coli* and in solution. Individual or combinations of different vectors carrying coding sequences of GFP–RAF40, His–SnRK2.4–FLAG, and His–HAB1–Myc, under *Ara*, *Lac*, and *Lac* operons, respectively, were transformed into *E. coli* BL21 (DE3) (Figure 6A). Strains were cultured in LB medium with 43 mM NaCl for 6 h and HAB1–Myc and His–SnRK2.4–FLAG were induced by IPTG for 2 h. Arabinose was then added to induce GFP–RAF40 expression for 1 h (Figure 6A). Aliquots were incubated ± 300 mM NaCl for 5 min, and Ser166 phosphorylation was determined in the lysate to indicate SnRK2.4 and RAF40 kinase activities. Because 800 mM mannitol is near its solubility limit, we instead used NaCl for *E. coli* treatment, as it provides a more convenient option for rapid stress application and activity assays. 1,6-HD was added as a control to abolish condensation. Wild-type SnRK2.4^WT^–FLAG has robust kinase activity as indicated by autophosphorylation (Figure 6B, lane 2). Co-expression of HAB1–Myc abolished the phosphorylation of SnRK2.4^WT^ (Figure 6B, lane 3). Inducing GFP–RAF40 expression with arabinose partially rescued SnRK2.4^WT^ activity in the strain co-expressing all three proteins (Figure 6B, lane 5). To better observe the effect of GFP–RAF40 on SnRK2.4 activation, we used the ‘kinase-dead’ isoform SnRK2.4^D131A^ (SnRK2.4^DA^), and detected Ser166 phosphorylation only in the strain co-expressing all three proteins (Figure 6B, lane 10). Five minutes of NaCl treatment dramatically enhances the phosphorylation of Ser166 in SnRK2.4^DA^ in five individual clones co-expressing three vectors (Figure 6C, lane 8 compared to lane 7, and S10A). As expected, 1,6-HD strongly down-regulates NaCl-triggered Ser166 phosphorylation (Figure 6C, lane 9), and results from four other individual clones replicate this (Figure S10B).

**Figure 6.**
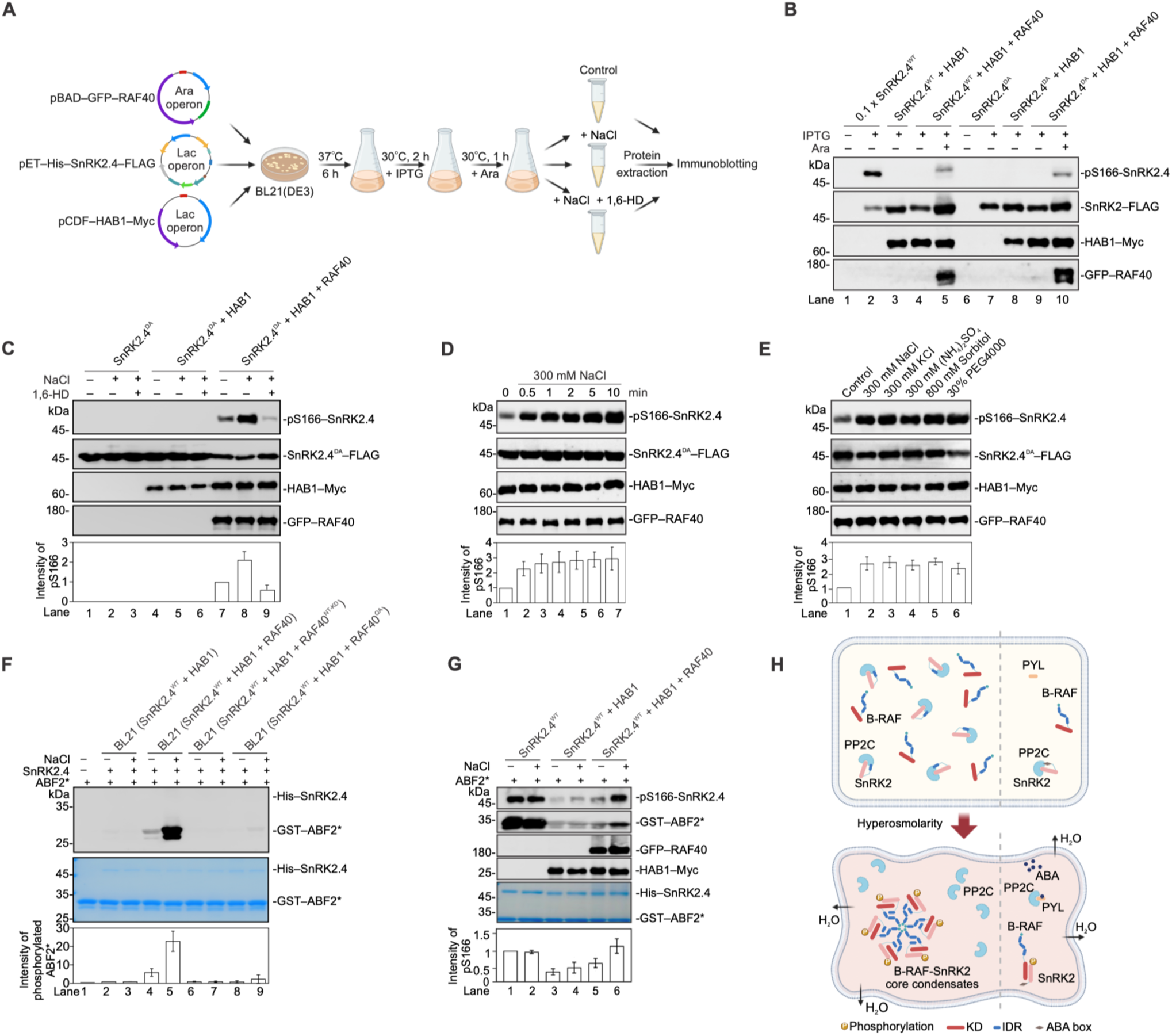
Reconstitution of the RAF–SnRK2 osmosensing core module in *E. coli* and *in vitro*. (**A**) Schematic of the experiment reconstituting the Arabidopsis osmosensing core module in *E. coli*. Briefly, the expression vector containing *SnRK2.4*, *HAB1* or *RAF40* under the *Lac* or *Ara* operon was transformed into BL21 (DE3), and screened with kanamycin, carbenicillin or spectinomycin resistance on solid low salt LB medium (containing 43 mM NaCl). The clones were cultured in liquid low salt LB medium and induced with 0.3 mM IPTG or 0.02% arabinose to express SnRK2.4, HAB1, or RAF40 proteins. Aliquots of culture were treated with NaCl or combined with 5% 1,6-HD and collected to extract total protein. The activation of RAF40 under NaCl-induced osmotic stress was assayed by the phosphorylation of SnRK2 (pS166-SnRK2.4) by immunoblotting. Anti–FLAG/Myc/GFP was used to probe the abundance of tagged SnRK2.4, HAB1, and RAF40. (**B**) Immunoblots of pS166-SnRK2.4, SnRK2.4^WT,^ or SnRK2.4^DA^–FLAG, HAB1-Myc, and GFP–RAF40 abundance in *E. coli* strains with different vector combinations. One-tenth of the aliquot was loaded in lane 2 to avoid over-exposure of pS166-SnRK2.4. (**C**) Immunoblots of pS166-SnRK2.4, SnRK2.4^DA^–FLAG, HAB1–Myc, and GFP–RAF40 abundance in *E. coli* strains with different vector combinations under ± 300 mM NaCl or 5% 1,6-HD treatment for 5 min. Relative intensity of pS166-SnRK2.4 (lower panel) was normalized with the bands in the control (lane 7) using ImageJ. Data represent mean ± SD (n = 5). (**D, E**) Immunoblots of pS166-SnRK2.4, SnRK2.4^DA^–FLAG, HAB1–Myc, and GFP–RAF40 abundance in *E. coli* strains expressing SnRK2.4^DA^–FLAG, HAB1–Myc, and GFP–RAF40 that treated with 300 mM NaCl for the indicated times (**D**), or the indicated chemicals for 5 min (**E**). Relative intensity of pS166-SnRK2.4 was normalized with the bands in the control (lane 1). Data represent mean ± SD (n = 3). (**G**) Thio-phosphorylation of ABF2 fragment by SnRK2.4 determined by *in vitro* phosphorylation assay in the presence of ATP-γS. His–SnRK2.4 was purified from *E. coli* strains carrying the indicated vectors, which were treated with ± 300 mM NaCl for 5 min before harvest. Recombinant GST–ABF2* (73–119 aa) was used as substrate to detect the kinase activity of SnRK2.4. Coomassie staining shows the abundance of His–SnRK2.4 and GST–ABF2*. Relative intensity of thio-phosphorylated ABF2 fragment was normalized with the bands in the SnRK2.4–HAB1 control (lane 2). Data represent mean ± SD (n = 3). (**H**) Phosphorylation and kinase activity of SnRK2.4 in the presence of HAB1 and RAF40 with NaCl treatment in an *in vitro* phosphorylation assay. His–SnRK2.4^WT^–FLAG, HAB1–Myc–His, His–SUMO–GFP–RAF40 were separately purified from *E. coli*. SnRK2.4 was incubated with HAB1 for 10 min, then reacted with RAF40 and 300 mM NaCl for 10 min in the presence of ATP-γS. ABF2 was used as substrate to detect the kinase activity of SnRK2.4. Immunoblots and Coomassie show thio-phosphorylated ABF2, pS166-SnRK2.4, HAB1–Myc, GFP–RAF40, SnRK2.4, and GST–ABF2*. Relative intensity of pS166-SnRK2.4 was normalized with the bands in the control of SnRK2.4 alone (lane 1). Data represent mean ± SD (n = 3). (**I**) Proposed model showing B4-RAF–SnRK2-mediated osmosensing of cellular hyperosmolarity in plants. Upon hyperosmolarity, B4-RAFs and SnRK2s form condensates together, spatially separated from the uncondensed PP2Cs, which mimics PYL-mediated inhibition of PP2C in ABA signaling. It promotes B4-RAF phosphorylation of SnRK2s to initiate osmotic-stress signaling. See also Figure S10–S12.

Phosphorylation of Ser166 in SnRK2.4^DA^ was observed after 30 s of 300 mM NaCl treatment (Figure 6D, lane 2). Similar to NaCl, application of KCl, ammonium sulfate, sorbitol, mannitol, or PEG4000 also strongly activates SnRK2.4^DA^ phosphorylation in *E. coli* strain co-expressing three proteins (Figure 6E). Deleting the IDRs, which abolish GFP–RAF40 condensation (Figure 2B and 3A), strongly impairs the ability of RAF40 to sense the hyperosmotic stress treatment and phosphorylate/activate SnRK2.4 (Figure S10C). Differing from the earlier observation that RAF40^IDR–KD^ cannot rescue the osmotic-stress hypersensitivity of *OK^1^*^30^*-null* (Figure 2E), co-expression of RAF40^IDR–KD^ slightly induces SnRK2.4^DA^ phosphorylation (Figure S10C, lane 4 compared to 3), suggesting that the NT of RAF40 has additional function *in planta*. Similar to RAF40, RAF24 also can sense hyperosmolarity and trigger SnRK2.4^DA^ phosphorylation in the *E. coli* system (Figure S10D).

We next examined the specificity of the IDR in B4-RAFs for osmosensing by substituting the IDR of RAF40 with that from other potential osmosensors. Replacing the RAF40 IDR with that of DCP5 (aa 92–301) failed to condense upon mannitol treatment *in vitro* (Figures S11A and S11B) and was unable to trigger SnRK2.4^DA^ phosphorylation in the *E. coli* reconstitution system (Figure S11C, lanes 4–6). Expression of the IDR^SEU^–RAF40 fusion protein (Figures S11D and S11E), in which the IDR of SEU (aa 1–308) was substituted, only weakly triggered SnRK2.4^DA^ phosphorylation in *E. coli* (Figure S11C, lanes 7–9). These results suggest a unique role of the B4-RAF IDR in hyperosmolarity sensing.

We then co-expressed the wild-type SnRK2.4 (SnRK2.4^WT^) in the reconstitution system and detected the activation of SnRK2.4^WT^ in the context of the ability to phosphorylate an ABF2 fragment (73–119 aa), a widely used SnRK2 substrate. ^4,49,50^ After incubation with IPTG and arabinose, His–SnRK2.4^WT^ was purified from culture aliquots without or with treatment of 300 mM NaCl for 5 min and incubated with the recombinant ABF2 fragment in the presence of ATP-γS as thiophosphate donor. As a result, purified His–SnRK2.4^WT^ from the strain co-expressing RAF40 shows stronger kinase activity than that from the strain without RAF40 (Figure 6F, lane 4 compared to 2). NaCl treatment could further increase the kinase activity, compared to that without NaCl treatment (Fold change = 21, *p* < 0.001, n = 4 independent replicates) (Figure 6F, lane 5 compared to 3). SnRK2.4^WT^ purified from the strain co-expressing RAF40^QA^ failed to phosphorylate ABF2 fragment *in vitro* (Figure 6F, lanes 8 and 9).

Besides B4-RAFs, certain protein kinases, such as BIK1^50^ and CPKs,^51^ may phosphorylate SnRK2s and participate in their activation under stress conditions. However, none of them could detectably trigger Ser166 phosphorylation of SnRK2.4 in response to NaCl treatment, when co-expressed with SnRK2.4 and HAB1 in the same *E. coli* system (Figure S11F). A member of the B2-RAF subgroup, RAF12, forms condensates upon NaCl treatment, and phosphorylates HAI2, an A-clade PP2C, to inhibit their activity.^52^ However, RAF12 also fails to detectably trigger phosphorylation of recombinant SnRK2.4^DA^ in *E. coli* in the presence of HAI2 (Figure S11G). Like SnRK2.4, the condensation of RAF12–GFP was also abolished by dysfunctions of B4 RAFs in *RAF12_pro_:RAF12–GFP/OK^1^*^30^*-null* plants (Figures S11H and S11I), indicating RAF12 alone may not be sufficient to sense hyperosmolarity and activate SnRK2.4, and the condensation/function of RAF12 may largely depend on B4-RAFs.

Last, we reconstituted the RAF40–SnRK2.4–HAB1 osmotic-stress sensing and signaling pathway in solution using purified proteins, with phosphorylation of the ABF2 fragment as the readout. Recombinant full-length SnRK2.4^WT^ displayed strong kinase activity, as indicated by both auto-thiophosphorylation and thiophosphorylation of the GST–ABF2 fragment (Figure 6G, lanes 1 and 2). Addition of recombinant HAB1 abolished SnRK2.4 kinase activity (Figure 6G, lanes 3 and 4). In the presence of HAB1, adding RAF40 alone only slightly restored SnRK2.4 activity (Figure 6G, lane 5). Supplementing NaCl to a final concentration of 300 mM, which triggered co-condensation of RAF40 and SnRK2.4, strongly reactivated SnRK2.4, as shown by robust thiophosphorylation of both SnRK2.4^WT^ and the ABF2 fragment (Figure 6G, lane 6). Although SnRK2.4 alone can form condensates *in vitro*, its reactivation still requires the presence of RAF40 (Figure 6G, lane 6 compared with lane 4).

### The B-RAF-mediated osmosensing module is conserved in plant and animal cells

The IDR is present in mammalian B-Raf and in plant B-subgroup RAFs, raising the question of whether B-RAFs play a conserved role in osmosensing. We randomly selected one B4-type RAF in rice, OsMAPKKK39, to test its condensation behavior *in vivo*. Indeed, the OsMAPKKK39–mVenus driven by its native promoter, but not the *35S*-promoter-driven mVenus control, underwent condensation in response to mannitol treatment (Figure S12A). Similarly, application of 100 mM mannitol induced punctate structures for B-Raf in HEK293T cells and concomitant phosphorylation of ERK1/2 (Figures S12B and S12C ). Treatment with RAF709, a specific B-Raf inhibitor, abolished mannitol-induced ERK1/2 phosphorylation, suggesting a potential role of B-Raf in osmosensing in human cells.

## DISCUSSION

Activation of the RAF–SnRK2 cascade is one of the earliest known events in osmotic-stress signaling that orchestrates most adaptation responses in plants. Here, we show that B4-subgroup RAFs can directly sense ionic and non-ionic hyperosmolarity through reversible condensation and form a signal hub with SnRK2s to relay osmotic-stress signals, independent of known osmosensors OSCA1^34^ and DCP5.^41^ Drought, high salinity, or low temperatures trigger fast water outflow or reduced water mobility in cells. The IDRs in B4-subgroup RAFs then directly sense hyperosmolarity in the cytosol and recruit SnRK2s to form co-condensates. In these core condensates, SnRK2s could be spatially separated from PP2Cs and phosphorylated/activated by B-RAFs, mimicking the protein–protein-interaction-based inhibition of PP2C in ABA signaling (Figure 6H). Activated SnRK2s then phosphorylate downstream effectors locally or relocate to other organelles to activate downstream adaptational events^19–21^, including mRNA processing^5^, chromatin remodeling,^53^ Ca^2+^ influx,^17^ vesicle formation,^54,55^ and stomatal closure. The basal kinase activity of B-RAFs is sufficient for SnRK2 activation *in vitro* and *in planta*;^11,23,30^ however, B-RAFs undergo fast hyperphosphorylation upon osmotic stimuli in plants.^4,38^ Besides autophosphorylation, we propose that some unidentified kinase(s) or partner(s) may integrate other osmotic signals from the cell wall, membrane, or cytosol to further enhance RAF activity via phosphorylation. Supporting this notion, our companied manuscript would be meritorious to identify the kinase(s)/partner(s) regulating condensation or activity of B-RAFs and to study crosstalk between RAF–SnRK2 core sensory condensates and other reported osmosensors, e.g., OSCA1 and DCP5. Our study suggests that co-condensation of RAF40 and SnRK2.4 is strictly controlled, and multiple regions in both proteins are essential. It would be interesting to elucidate the cue that determines the mechanism and identify the additional components involved in formation of RAF–SnRK2 core condensates *in planta*. B-RAF–SnRK2 co-condensation occurs upon both high concentrations of ions (NaCl, KCl, (NH_4_)_2_SO_4_) and low-molecular-weight chemicals (PEG, mannitol, sorbitol), and macromolecular electrolytes (PEG), while DCP5 and SEUSS only respond to PEG but not NaCl. ^39,41^ We propose that IDRs in the B-RAF–SnRK2 core module can sense water-potential changes caused by these hyperosmotic agents, distinct from IDRs in DCP5 or other potential osmosensors. Mutation of the spacer residues, polar uncharged glutamine to alanine abolishes RAF40 condensation, supporting this notion. These changes could be a wide range of physical stimuli, influencing molecular dipole–dipole, dipole-induced dipole, hydrogen bonding, ion–dipole, and ion–ion interactions, spectroscopic/thermodynamic changes, thus causing phase separation due to interfacial tension, ^56,57^ which is of interest to explore.

Roles for LLPS or condensation have been established in sensing physical environmental stimuli such as hydration in seed germination,^58^ cold,^59^ or high temperature^60–63^ in plants. DCP5 senses cell-volume changes triggered by hyperosmolarity.^41^ Most of these reported sensors, including DCP5, are transcriptional regulators. Among them, FLOE1,^58^ RBGD2/4,^62^ TWA1,^63^ SEUSS,^39^ and DCP5^41^ are activators, while FRIGIDA^59^ and EFL3^60^ are co-suppressors. Different from these transcriptional regulators, B-RAFs and SnRK2s are the central signaling proteins that together orchestrate the multifaceted cellular adaptive responses to osmotic stress. These kinase pairs can sense hyperosmotic stress and form co-condensates, physically evading the inhibition by the PP2C protein phosphatases; B-RAF then phosphorylates and activates SnRK2s in the co-condensates. This efficient and straightforward osmosensing and signal relaying by this core module can be reconstituted in *E. coli* and *in vitro* in a test tube, and in the presence of the inhibitory phosphatase HAB1 (Figure 5B). Mammalian B-Raf also forms condensates and participates in osmotic-stress-triggered ERK1/2 phosphorylation. Thus, we propose these spatially separated core condensates of active enzymes forming away from inhibitors, or *vice versa*, could be a general strategy for forming signal hubs for plants (and potentially other organisms, e.g. WNK1 signaling in animals ^64^) to rapidly respond to environmental and endogenous stimuli.

### Limitations of the study

In this study, we demonstrate that both ionic and non-ionic hyperosmolarity rapidly trigger the condensation of B4-RAFs. Mutation of polar, uncharged glutamine residues in the IDR abolishes RAF40 condensation and the subsequent activation of SnRK2s. Due to current technical limitations, we could not directly visualize the physical changes of B4-RAFs, particularly within their IDRs, under osmotic stress. Moreover, individual B4-RAF members exhibit distinct sensitivities to hyperosmolarity, but how sequence variation in IDRs relates to osmosensitivity remains unclear. Beyond the IDR, other domains also contribute to B4-RAF condensation and activation, yet their precise functions in osmosensing and relaying remain to be elucidated.

In Arabidopsis, more than 20 B-RAFs and 10 SnRK2s constitute a complex kinase network to perceive and relay hyperosmotic stress. We identify B4-RAFs as the central osmosensors of this cascade: in the *OK^1^*^30^*-null* mutant, both condensation and activation of SnRK2.4 are abolished. Strikingly, RAF12 condensation is also impaired in *OK^1^*^30^*-null*, suggesting that B4-RAFs may promote the activation of B2 and B3 RAFs through mechanisms that are not yet understood. How such a multilayered kinase cascade is organized and activated within minutes of osmotic stress remains an open question.

Some other osmosensors, such as OSCA1 and DCP5, have been reported in plants. Although our data indicate that B4-RAFs act independently of OSCAs and DCP5, it is plausible that plant cells integrate osmotic cues from multiple osmosensors located in different organelles. Understanding this integration represents an important future challenge. Similarly, in mammalian cells, how osmosignals perceived by B-Raf and WNK1 are coordinated to regulate downstream events also remains unresolved. The precise role of B-Raf in osmosensing and its contribution to mammalian stress signaling will require detailed investigation.

## RESOURCE AVAILABILITY

### Lead contact

Further information and requests for resources and reagents should be directed to and will be fulfilled by the lead contact, Pengcheng Wang.

### Materials availability

This study did not generate new, unique reagents. The plant materials and plasmids generated in this study should be directed to and will be fulfilled by the lead contact.

### Data and code availability

Any additional information required to reanalyze the data reported in this paper is available from the lead contact upon request.

## ACKNOWLEDGMENTS

This work was supported by the National Key Research and Development Program of China, Grant 2021YFA1300402 (to P.W.). We are grateful to Drs. Zhizhong Gong of China Agriculture University, Siyi Guo of Henan University, and Tongda Xu of Fujian Agriculture and Forestry University for providing the ABI1, HAB1, and ABI2 fluorescent lines, Hongwei Guo of Southern University of Science and Technology for providing the *dcp5* mutant, Xiaofeng Fang of Tsinghua University for providing the SEUSS vector, and Jiamu Du, Jun Tian, and Wen Zhou of Southern University of Science and Technology for helpful discussions.

## AUTHOR CONTRIBUTIONS

P.W. designed experiments, acquired funding, performed data analysis, and wrote the paper. J.K.Z., C.P.S., and L.Z. participated in data analysis and wrote the paper, G.Liu., Z.L., G.Lin., X.W., and X.L. designed and performed experiments, and performed data analysis.

## SUPPLEMENTAL INFORMATION

Document S1. Figures S1–S12.

Video S1. Time-lapse imaging showing the recovery after bleaching of RAF40–GFP condensates *in vitro*, related to Figure 1.

Tables S1. Primers used in this study.

## STAR★METHODS

### KEY RESOURCES TABLE

#### EXPERIMENTAL MODEL AND STUDY PARTICIPANT DETAILS

##### Plant materials and growth conditions

*Arabidopsis thaliana* (wild-type Col-0), *OK^130^-null*,^4^ *35S_pro_:GFP–RAF11* /Col-0,^4^ *35S_pro_:GFP–RAF20* /Col-0,^4^ *35S_pro_:ABI2–GFP*/Col-0,^46^ *HAB1_pro_:HAB1–Venus*/Col-0,^47^ and *ABI1_pro_:ABI1–GFP*/*abi1 abi2* ^48^ transgenic lines were previously described.

To generate the full-length and truncated *RAF40_pro_:RAF40s–GFP–FLAG*/*OK^1^*^30^*-null* and *RAF40_pro_:RAF40–GFP–FLAG*/*snrk2-sep* transgenic lines, the promoter (2 kb) and coding sequences of *RAF40* full-length or truncated coding sequences of *RAF40* (*RAF40^IDR–KD^*, *RAF40^NT–KD^*, *RAF40^NT–IDR^*, *RAF40^NT^, RAF40^IDR^*, or *RAF40^KD^* variants) and *RAF40^QA^* were amplified and cloned into the binary vector pCAMBIA2300–GFP–FLAG and transformed into the *OK^1^*^30^*-null* mutant or *snrk2-sep* (*snrk2.1/4/5/7/8/9/10*) mutant.^22^ Similarly, *RAF24* (1.4 kb promoter and full-length genomic sequence) and *RAF8* (2 kb promoter and coding sequence) were cloned into pCAMBIA2300–GFP–FLAG and transformed into *OK^1^*^30^*-null* or Col-0, respectively. For *SnRK2.4* lines, a 1.1 kb promoter and coding sequence were fused to GFP or RFP or BFP in pCAMBIA2300-FLAG and introduced into *OK^130^-null* or Col-0. Immunoblots with corresponding antibodies were used to determine the presence and amount of transgenic protein to avoid over-expression in the positive lines.

To generate *35S_pro_:RAF40–GFP SnRK2.4_pro_:SnRK2.4–RFP* transgenic Arabidopsis lines, the multigene construct containing *35S_pro_:RAF40–GFP* and *SnRK2.4_pro_:SnRK2.4–RFP* was assembled using the UNiE-mediated DNA assembly (UNiEDA) strategy as previously described.^65^ Briefly, gene-expression cassettes were amplified with chimeric primers containing unique nucleotide sequences (UNSs) and digested with Nb.BtsI. The digested fragments were ligated with the Nb.BtsI-treated binary vector pYL1300H-UNiE in a one-step thermal-cycling ligation reaction. Verified plasmids were introduced into *A. tumefaciens* GV3101 and transformed Col-0, *OK^1^*^30^*-null,* or *snrk2-sep* Arabidopsis plants.

Seeds of Arabidopsis thaliana (wild-type Col-0), mutant, and transgenic lines were surface-sterilized with 75% (v/v) ethanol for 2 min and rinsed five times with sterile water. Sterilized seeds were sown on half-strength Murashige & Skoog (1/2 MS) medium plates, cold-stratified at 4°C for 2 d, and subsequently transferred to a growth chamber maintained at 21°C with 50–60% humidity under a 16/8 h light/dark cycles (100 μmol photons m^−2^ s^−1^). After germination, seedlings were moved to pots containing a soil mixture of vermiculite and garden soil (1:4, v/v) and grown in a greenhouse under the same environmental conditions as the growth chamber.

Tobacco (*Nicotiana benthamiana*) was grown in the cultivating room at a photoperiod of 16 h light/8 h dark with the temperature at 24°C, 70% humidity and the light intensity of 140 μmol m^−2^ s^−1^, and normally 4-week-old tobacco seedlings were used for transient expression of the proteins.

For transgenic rice plants, the promoter (1.8 kb) and coding sequences of *OsMAPKKK39* (LOC_Os06g08280) was cloned into the pCAMBIA1300–mVenus vector to generate the *OsMAPKKK39_pro_:OsMAPKKK39–mVenus* construct. The resulting plasmids were introduced into *A. tumefaciens* EHA105 and then transformed into the calli of *Oryza sativa* cv. Nipponbare. For stress treatment, rice was grown in a growth room (14 h light at 30°C/10 h dark at 25°C) with a relative humidity around 60%.

#### METHOD DETAILS

##### Recombinant protein expression and purification

For recombinant protein expression in *E. coli*, full-length *RAF40*, *RAF24* and *RAF20* were cloned into the pET–10×His–SUMO–GFP with an N-terminal 10×His*–*SUMO and GFP tag. The vectors for RAF40 and variants, RAF40^NT^, RAF40^IDR^, RAF40^KD^, RAF40^IDR–KD^, RAF40^NT–KD^,RAF40^NT–IDR^, RAF40^QA^, RAF40^YF^, and RAF40^RK^ were generated from RAF40 vectors. The full-length coding sequences of *SnRK2s*, *HAB1, HAB2, PP2CA* were cloned into the pET28a vector with N-terminal 6×His tag and C-terminal YFP or RFP tag, respectively.

Full-length RAF40, RAF24, RAF20 and their variants were expressed in *E. coli* at 18°C for 16 h with 0.3 mM IPTG induction (low salt LB containing 2.5 g/L NaCl for RAF24 and RAF20). Cells were lysed in buffer [50 mM Tris-HCl, pH 7.5, 100 or 50 mM (for RAF24 and RAF20) NaCl, 50 mM imidazole, 10% v/v glycerol, and 2 mM PMSF], and the supernatant was loaded onto a Ni-affinity column (GE Healthcare, USA). Bound proteins were eluted with buffer containing 500 mM imidazole, and the 10×His–SUMO tag was removed by His–Ulp1 protease during dialysis [50 mM Tris-HCl, pH 7.5, 100 mM or 20 mM (for RAF24 and RAF20) NaCl] at 4°C for 10 h. The mixture was passed through a second Ni column to remove the cleaved tag and uncleaved proteins.

##### Plant image acquisition

Roots of 8-d-old Arabidopsis seedlings and rice plants grown on 1/2 MS medium, for osmotic stress treatments, Arabidopsis seedlings and rice seedlings were incubated in 1/2 MS liquid medium without or with indicated concentration mannitol for the indicated times. prior to imaging. Arabidopsis seedlings subjected to temperature stress were treated at 4 °C or 37 °C and subsequently imaged. Samples were then transferred onto a slide and imaged on a Zeiss LSM880 confocal laser microscope (details below).

For RAF40 condensate-reversibility assays, roots of 8-d-old Arabidopsis seedlings were soaked in 1/2 MS liquid medium with 800 mM mannitol as described above, and after imaging, mannitol was washed off from the roots with 1/2 MS liquid medium, and the same roots were imaged in 1/2 MS liquid medium after 10 min. Roots were transferred into 1/2 MS liquid medium with 800 mM mannitol again for 2 min and imaged to assess their ability to re-form condensates.

To measure the co-localization of RAF40 and SnRK2.4 *in vivo*, the *SnRK2.4_pro_:SnRK2.4–RFP* construct was introduced into *RAF40_pro_:RAF40–GFP/OK^130^-null* plants by *Agrobacterium*-mediated transformation, and immunoblots with anti-GFP were used to determine the presence and amount of transgenic SnRK2.4–RFP. Confocal imaging was conducted on the root tips.

##### Live-cell imaging of B-Raf condensates in HEK293T cells

HEK293T cells were plated in 6-well dishes in complete media and allowed to grow 80*–*90% confluency. Cells were transfected in antibiotic-free media the following day with standardized amounts of GFP-tagged B-Raf plasmids. Cells were re-plated 24h post-transfection on poly-D-lysine-coated Biosharp confocal dishes. Tonic solutions and media were prepared and warmed to 37°C on the day of imaging.

##### Confocal laser-scanning microscopy

Unless otherwise indicated, all imaging was done with the Zeiss 880 Airyscan inverted confocal laser-scanning microscope using a 40×/1.2 water correction objective. BFP, GFP (eGFP), YFP/mVenus, and mCherry/RFP were excited at 399, 488, 514, and 561 nm, respectively, and emission was collected at 445–454, 500–520, 525–555, and 575–615 nm using appropriate band-pass filters. When eGFP/YFP were imaged together with mCherry, a spectral GASP detector was used to collect emission from eGFP/YFP. For co-localization experiments, samples were imaged sequentially between each line to ensure that the co-localization signals were not due to bleed-throughs, and co-localization analysis was performed using Fiji (version J 1.54f).

For HEK293T cells, live-cell imaging was performed on a Leica STELLARIS 5 confocal laser-scanning microscope using the 40× objective with 488 nm excitation and 500–550 nm emission wavelengths, and representative images were captured. Deconvolution processing was applied to images of Arabidopsis roots and *in vitro* puncta to improve image clarity and resolution.

##### *In vitro* condensation assay

Purified proteins in dialysis buffer were diluted to indicated concentrations in reaction buffer (20 mM HEPES, pH 7.5, 50 mM NaCl, and centrifuged at 16,000 × *g* at 4°C for 10 min to remove protein aggregates. Proteins were mixed with the indicated chemicals including mannitol, sorbitol, erythrose, LiCl, (NH_4_)_2_SO_4_, KCl, NaCl, ethylene glycol (EG), PEG200, PEG1000, PEG4000, PEG600, PEG8000, individually, to the indicated concentration, and incubated for the indicated times before confocal imaging.

For co-condensation assays, SnRK2.1–YFP, SnRK2.4–YFP, SnRK2.5–YFP, SnRK2.9–YFP, or SnRK2.10–YFP, were mixed with HAB1–RFP, PP2CA–RFP, or HAB2–RFP gently in buffer containing 20 mM HEPES, pH 7.5, 50 mM NaCl, without or with 800 mM mannitol.

After incubation for 5 min, GFP–RAF40, GFP–RAF24, or GFP–RAF20 was then added and mixed gently. Images were captured within 5 min.

##### Condensation assay of GFP–RAF40 and GFP–RAF24 in *E. coli*

Full-length *RAF40* and *RAF20* coding sequences were cloned into the pBAD–GFP expression vector (Thermofisher, V43001). Proteins were expressed in *E. coli* BL21 (DE3) cells in the low salt LB medium (2.5 g/L NaCl) with presence 0.02% w/v arabinose for 3 h at 30°C. After induction, cells were collected and resuspended in fresh liquid low salt LB medium without or with 800 mM mannitol. Cells were immediately used for imaging on a Zeiss LSM880 confocal laser-scanning microscope.

##### Condensate measurement

All condensates were measured by FIJI. For droplet number and area, condensates were selected through the “Threshold” function by adjusting the brightness range. To refine the condensate segmentation, internal gaps were filled using the “Fill Holes” function, and overlapping structures were separated via the “Watershed” algorithm. Subsequently, noise within the selected regions was eliminated using the “Analyze Particles” function with a defined particle size threshold. The total number of remaining condensates and the area of each condensate were automatically calculated by applying the analysis.

For colocalization of droplets, Images were split by channels, background-subtracted, and thresholded to generate binary masks. Overlapping droplets were identified using “Image Calculator”. The colocalization ratio was calculated as the percentage of overlapping puncta relative to total puncta in channel. At least 10 cells from three independent experiments were quantified.

##### Protoplast preparation and transformation

Mesophyll protoplasts were isolated from 4-week-old Col-0 leaves using a modified protocol ^11^. Leaf strips were digested in enzyme solution (cellulase R-10 and macerozyme R-10, Yakult Pharmaceutical Industry) for 3 h in the dark, filtered, and centrifuged at 150 × *g* for 3 min at 4°C. Protoplasts were washed with W5 buffer (2 mM MES, pH 5.7, 154 mM NaCl, 125 mM CaCl_2_, 5 mM KCl), kept on ice for ≥ 30 min, and resuspended in MMg buffer (4 mM MES, pH 5.7, 0.4 M mannitol, 15 mM MgCl_2_). Fifty micrograms of plasmids were transfected using the PEG-calcium method. After 16 h in the dark, protoplasts were transferred to fresh MMg or MMg with additional 400 mM mannitol for 30 min before imaging.

##### Phenotyping of root length

For growth assays, surface-sterilized seeds were grown vertically on 1/2 MS, pH 5.7, 0.85% w/v agar and kept at 4°C for 2 d. Seedlings were grown vertically for 3–4 d and then transferred to medium with or without 200 mM mannitol or 100 mM NaCl. Root length was measured at the indicated number of days.

##### Reconstitution of the RAF–SnRK2 signaling module in *E. coli*

For co-expression of recombination proteins in *E. coli*, coding sequences of SnRK2.4–FLAG, HAB1/HAI2–Myc, and GFP–RAF40/RAF24/RAF12/BIK1/CPK6, were cloned into pET28a, pCDF (Merck, 71340), and pBAD vectors, respectively. Positive colonies were selected from low-salt LB medium with 5g/L NaCl containing the corresponding antibiotic and cultured in low-salt LB liquid medium at 37°C. When OD_600 nm_ reached 1.0, a final concentration of 0.3 mM IPTG was added to induce the expression of SnRK2.4*–*FLAG and HAB1/HAI2*–*Myc for 2 h at 30°C. A final concentration of 0.02% w/v arabinose was then added to induce the expression of GFP–RAF40/RAF24/RAF12/BIK1/CPK6 for 1 h at 30°C. The bacterial culture was aliquoted and treated with indicated concentration of chemicals without or with and 5% w/v 1,6-HD for the indicated times. For mannitol treatment, bacteria were collected by centrifugation and resuspended in medium with 800 mM mannitol, as the 800 mM mannitol is near its saturated solution (about 972 mM).

After treatment, cells were collected by centrifuged at centrifuged at 16,000 × *g* at 4°C, and pellets were lysed in extraction buffer (100 mM HEPES, pH 7.5, 5 mM EDTA, 5 mM EGTA, 10 mM DTT, 10 mM Na_3_VO_4_, 10 mM NaF, 50 mM β-glycerophosphate, 1 mM PMSF, 5 μg/mL leupeptin, 5 μg/mL antipain, 5 μg/mL aprotinin, and 5% glycerol) and denatured in SDS loading buffer at 95°C for 10 min with vigorous shaking (1,500 rpm). The proteins were separated by 8% SDS*–*PAGE and immunoblotted with the indicated antibodies. The phosphorylation of Ser166 in SnRK2.4 was determined using a custom antibody recognizing the conserved serine residue corresponding to Ser166 in SnRK2.4 ^24^. Abundance of SnRK2.4, PP2Cs, and RAF24/RAF40/RAF12/BIK1/CPK6 was detected using anti-FLAG, Anti-Myc, and anti-GFP antibodies, respectively. The list of antibodies is reported in the key resources table.

##### Preparation of cell lysates and immunoblot analysis

To examine the natively expressed B-Raf–ERK pathway under hyperosmotic stress, unedited HEK293T cells were grown in 6-well dishes. Once cells reached 90% confluence (∼24 h), cells were pretreated with the specific 50 nM B-Raf inhibitor RAF709 (SelleckChem, S8690) or DMSO for 2 h, followed by replacement of growth media with fresh media supplemented with the indicated concentration of mannitol for 15 min. Collected cells were washed with PBS (137 mM NaCl, 2.7 mM KCl, 10 mM Na_2_HPO_4_, 1.8 mM KH_2_PO_4_, passed through a 0.22-µm filter), and cytosolic proteins were extracted from cell pellets with ice-cold RIPA Lysis Buffer (50 mM Tris-HCl pH 7.4, 150 mM NaCl, 0.5% sodium deoxycholate, 0.1% w/v SDS, 10 mM NaF, 5 mM EDTA, 1% v/v Triton X-100, supplemented with protease and phosphatase inhibitors). After centrifugation, protein concentrations in the supernatant were determined using the Pierce BCA Protein Assay Kit (Thermo Fisher Scientific). 20-μg protein extracts were boiled in SDS loading buffer, separated by 10% SDS*–*PAGE and immunoblotted with the indicated antibodies. The phosphorylation of ERK was determined with anti-pERK1/2 antibody (CST, 4370S) and anti-GAPDH (Abmart, M20006F) was used as loading control.

#### QUANTIFICATION AND STATISTICAL ANALYSIS

Statistical parameters including the definitions and exact values of n (e.g., number of biological repeats, number of plants, number of cells, etc.), distributions, and deviations are reported in the Figures and corresponding Figure Legends. Comparisons between two groups were performed using two-tailed Student’s *t* test. For multiple comparisons, one-way ANOVA followed by Tukey’s post hoc test was used, and statistically significant differences among groups are indicated by letters (a–d) in the Figures. Significance is reported as *p* > 0.05 (not significant), ^∗^*p* < 0.05, ^∗∗^*p*< 0.01, ^∗∗∗^*p*< 0.001,^∗∗∗∗^*p* < 0.0001. All statistical analyses were performed using GraphPad Prism software following standard procedures.

