## Supplemental Figures for "Rapid sensing and relaying of cellular hyperosmotic-stress signals via RAF–SnRK2 core condensates"

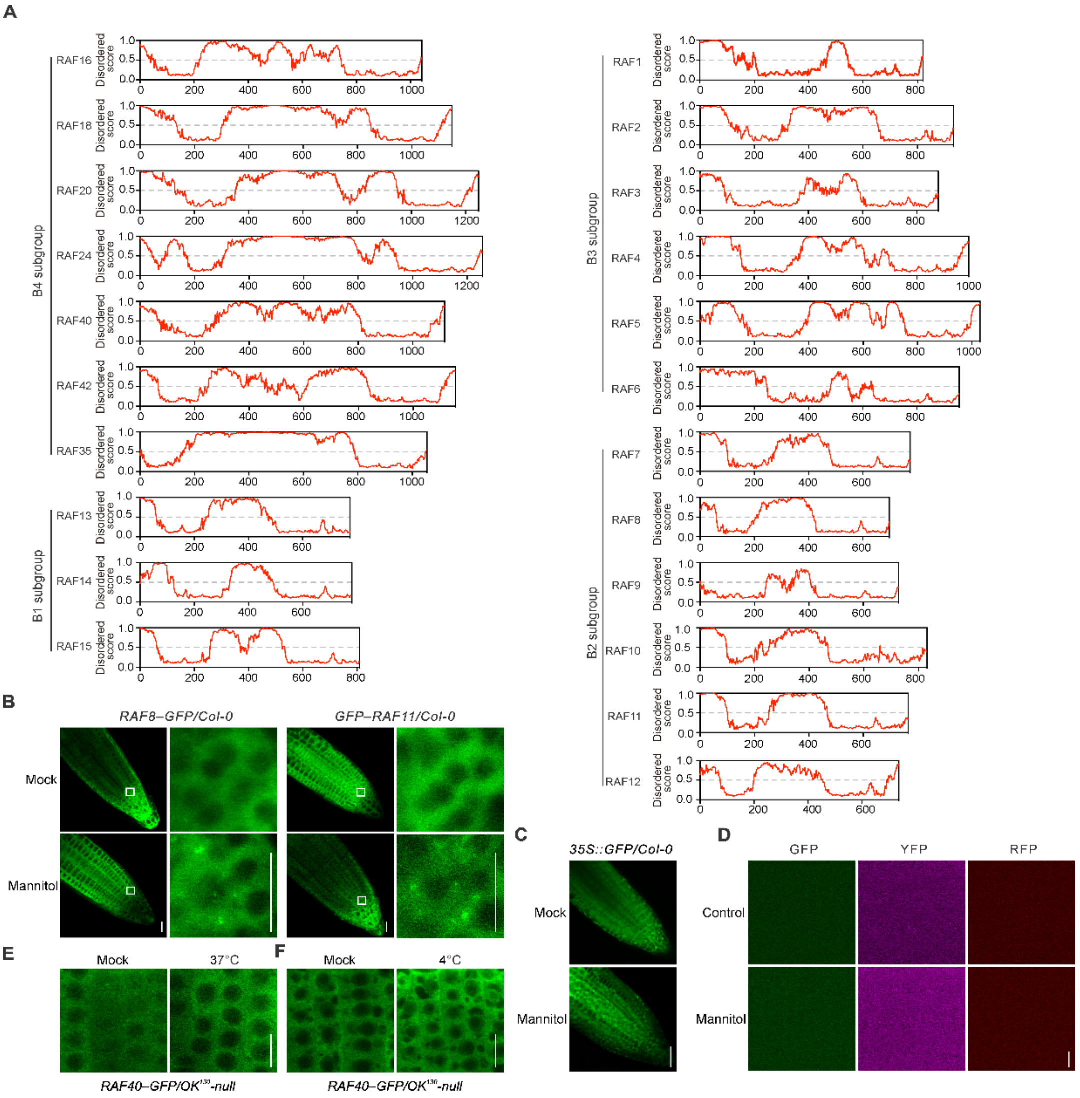


**Figure S1. B-RAFs are predicted to contain IDRs and B2-subgroup members RAF8 and RAF11 undergo mannitol-triggered condensation. Related to Figure 1.**

(A) The intrinsically disordered regions of subgroup 1–4 B-RAFs predicted using IUPred2.

(B) Mannitol treatment triggers RAF8–GFP and RAF11–GFP condensation in transgenic plants. The small white boxes in the left-hand images are zoomed in at right.

(C, D) Representative confocal microscopy images of free GFP *in planta* (*$35S_{pro}$:GFP*) (C) or recombinant GFP/YFP/RFP alone *in vitro* (D) with 800 mM mannitol.

(E, F) Heat and cold treatments do not trigger the condensation of RAF40–GFP *in vivo*.

Scale bars: 10 μm in (B), (C), (E) and (F); 2 μm in (D).

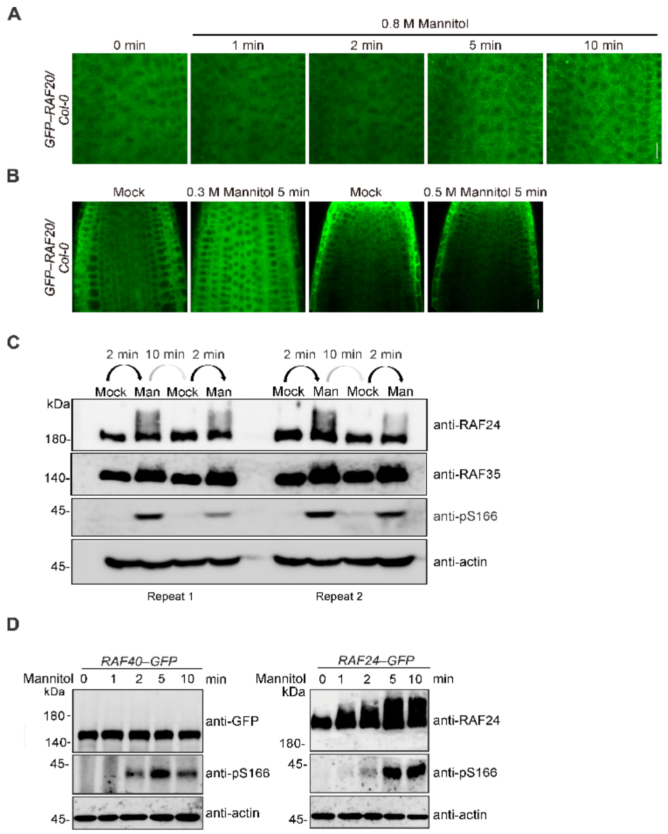


**Figure S2. Mannitol-triggered condensation and mobility shift of RAF20, RAF24, and RAF35. Related to Figure 1.**

(A) Distribution dynamics in root-tip cells of *GFP–RAF20*/Col-0 transgenic plants upon 800 mM mannitol treatment for the indicated times.

(B) Distribution dynamics of RAF20–GFP in *GFP–RAF20/*Col-0 transgenic plants in 1/2 MS (Mock) or 1/2 MS supplemented with 300 or 500 mM mannitol.

(C) Immunoblots of RAF24, RAF35, pS166-SnRK2.4 and actin abundance. Arabidopsis seedling was treated with 800 mM mannitol for 2 min, followed with recovery in 1/2 MS medium for 10 min, and then retreated 800 mM mannitol for 2 min.

(D) Immunoblots of RAF40–GFP, RAF24–GFP (top panel), Ser166 phosphorylation in SnRK2.4 (middle), and the loading control actin (bottom). Arabidopsis seedlings were treated with 800 mM mannitol for indicated time.

Scale bars: 10 µm in (A) and (B).

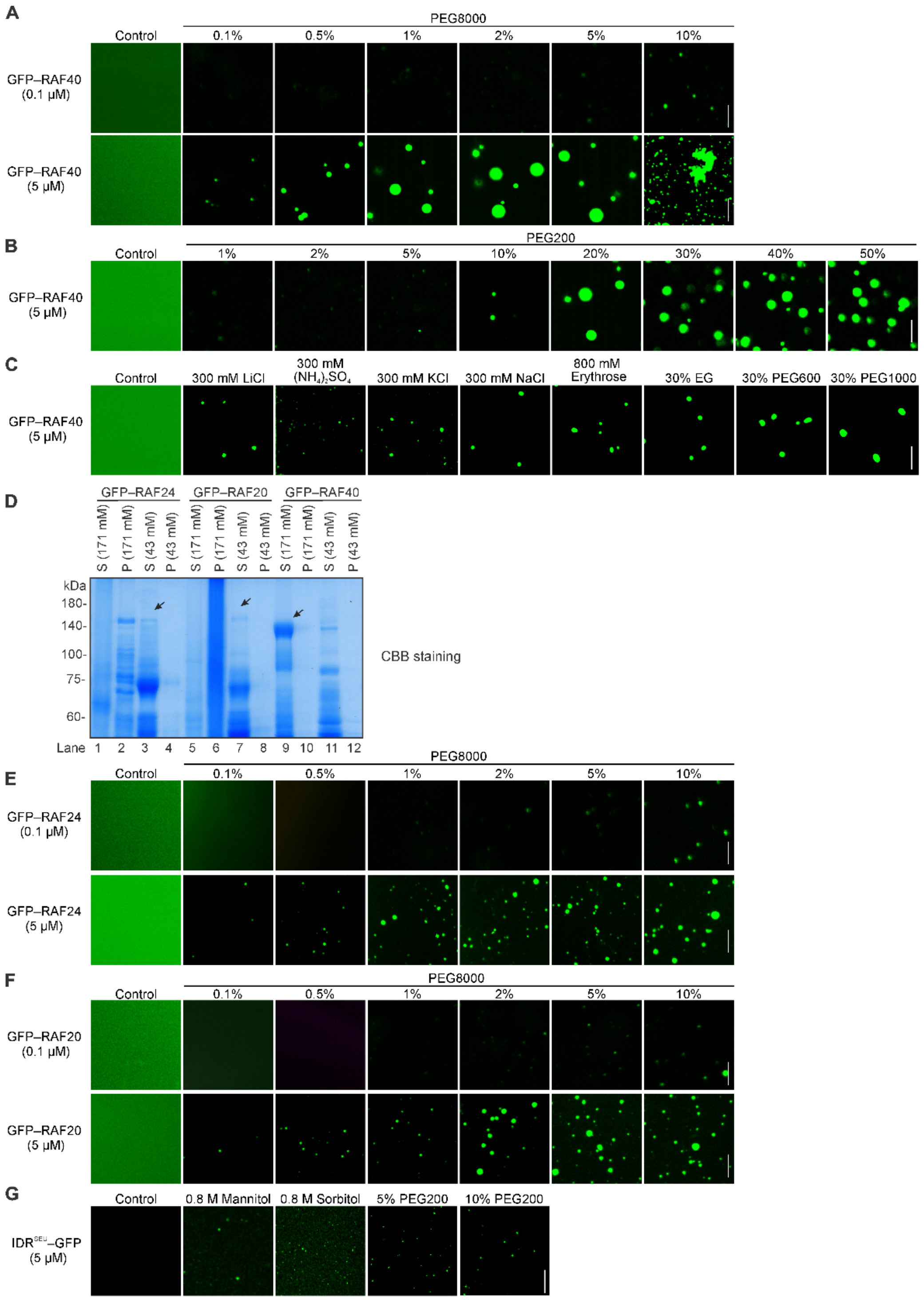


**Figure S3. Condensation of recombinant B4-RAFs in different hyperosmolarity treatments. Related to Figure 1.**

(A) Confocal micrographs of GFP–RAF40 condensates formed at the indicated protein and PEG8000 concentrations.

(B, C) Confocal micrographs of 5 μM GFP–RAF40 condensates formed at the indicated concentrations of PEG200 (B) and indicated concentrations of chemicals (C).

(D) Coomassie Brilliant Blue (CBB) staining of GFP–RAF24, GFP–RAF20, and GFP–RAF40

expression in LB culture media with 10 g/L (about 171 mM), or 2.5 g/L (about 43 mM) NaCl. S, supernatant; P, pellet.
(E, F) Confocal micrographs of GFP–RAF24 (E), and GFP–RAF20 (F) condensates formed at the indicated protein and PEG8000 concentrations.
(G) Confocal micrographs of 5 μM IDR$^{SEU}$–GFP condensates formed at the indicated concentrations of mannitol, sorbitol, and PEG200.
Scale bars: 5 μm in（A–C) and（E–G).

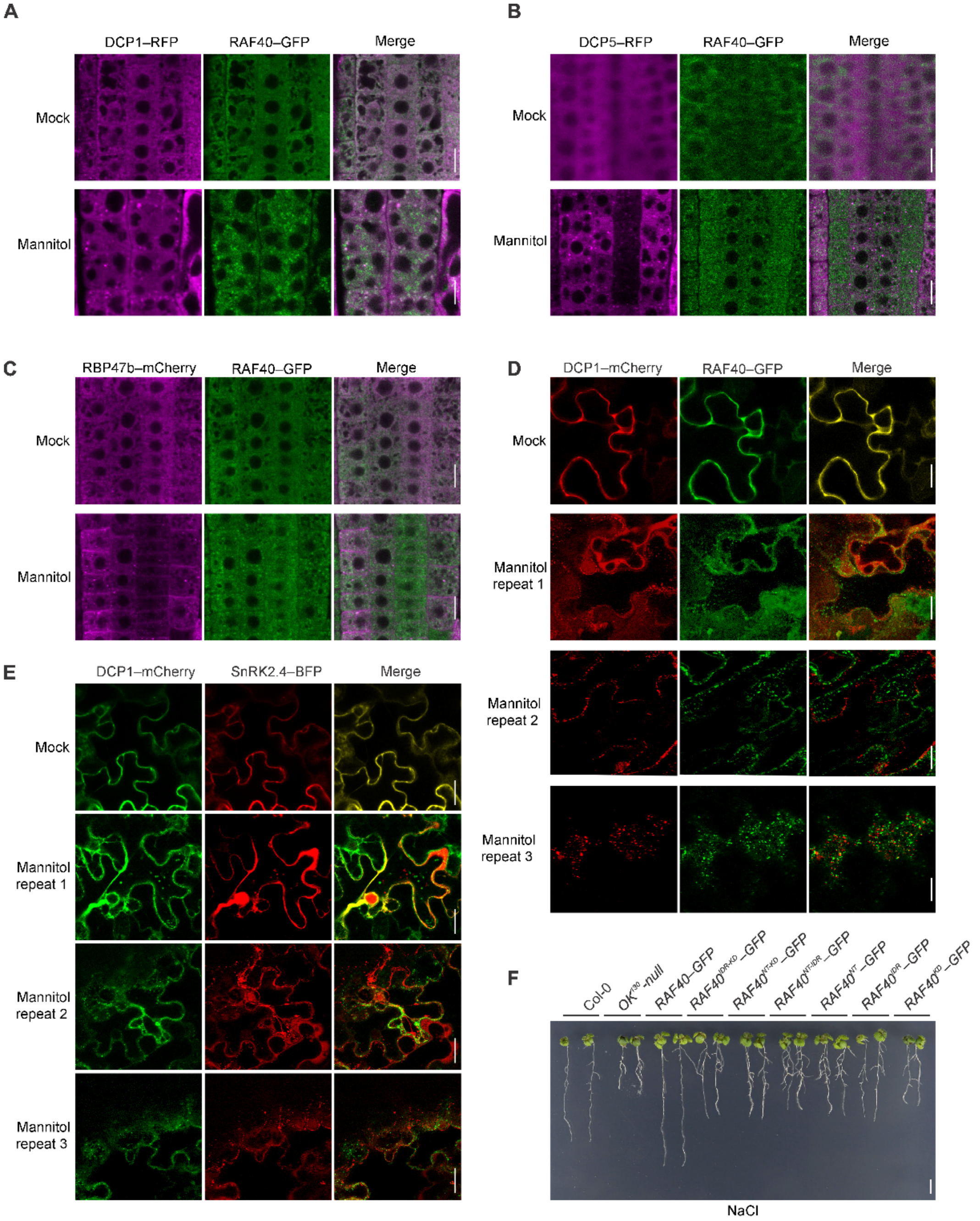


**Figure S4. RAF40–GFP puncta are distinct from DCP1, RBP47b, and DCP5 under osmotic stress and growth of *$OK^{130}$-null* complementation lines on NaCl. Related to Figure 2.**

(A–C) Confocal micrographs of transgenic Arabidopsis root cells expressing both RAF40–GFP and DCP1–RFP (A), DCP5–RFP (B), RBP47b–mCherry (C) treated with or without 800 mM mannitol for 5 min.

(D, E) Confocal micrographs of tobacco leave transiently co-expressing DCP1–mCherry and RAF40–GFP (D) and SnRK2.4–BFP (E).

(F) Photograph of 7-d-old seedlings of Col-0, *$OK^{130}$-null* and different *$OK^{130}$-null* complementation lines grown on 1/2 MS medium with 100 mM NaCl. Scale bar: 1 cm.

Scale bars: 10 µm in (A)–(E), 1 cm in (F).

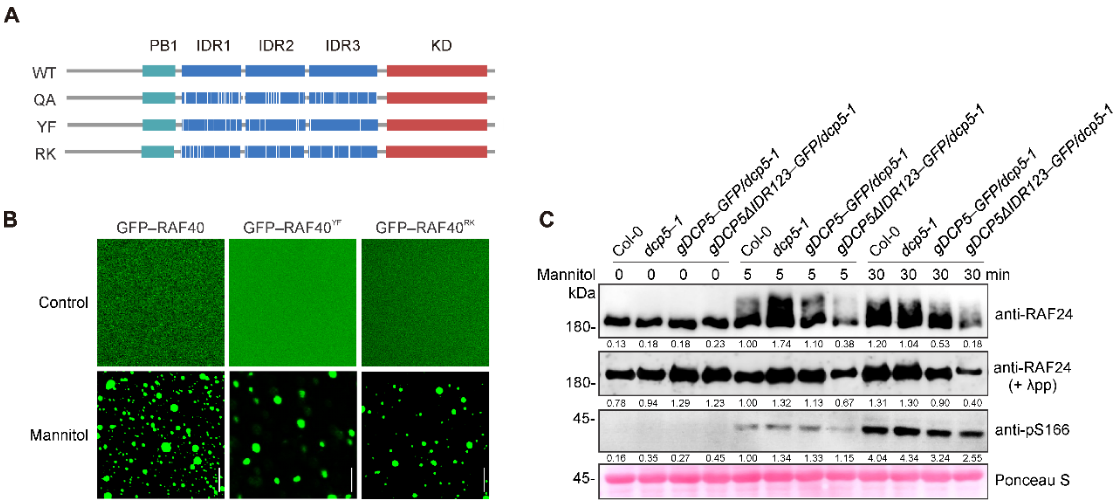


**Figure S5. Point mutations or dysfunction of DCP5 do not alter RAF40 condensation or activation. Related to Figure 3.**

(A) Schematic representation of the intrinsically disordered region (IDR) of RAF40, with the positions of 36 glutamine (Q), 15 tyrosine (Y) and 27 arginine (R) residues indicated by white vertical lines.

(B) Fluorescence images showing condensate formation of recombinant GFP–RAF40$^{YF}$ and GFP–RAF40$^{RK}$ variants. Scale bar: 5 µm.

(C) Immunoblots showing the mobility shift and abundance of RAF24 and pS166-SnRK2.4 in *dcp5* mutant or complementation lines. Seedlings of Col-0, *dcp5-1*, and its transgenic complementation lines were treated with 800 mM mannitol for 5 or 30 min. λPP phosphatase was used to dephosphorylation of total protein (second panel) to better show the abundance of native RAF24. Ponceau S was used as a loading control (bottom panel).

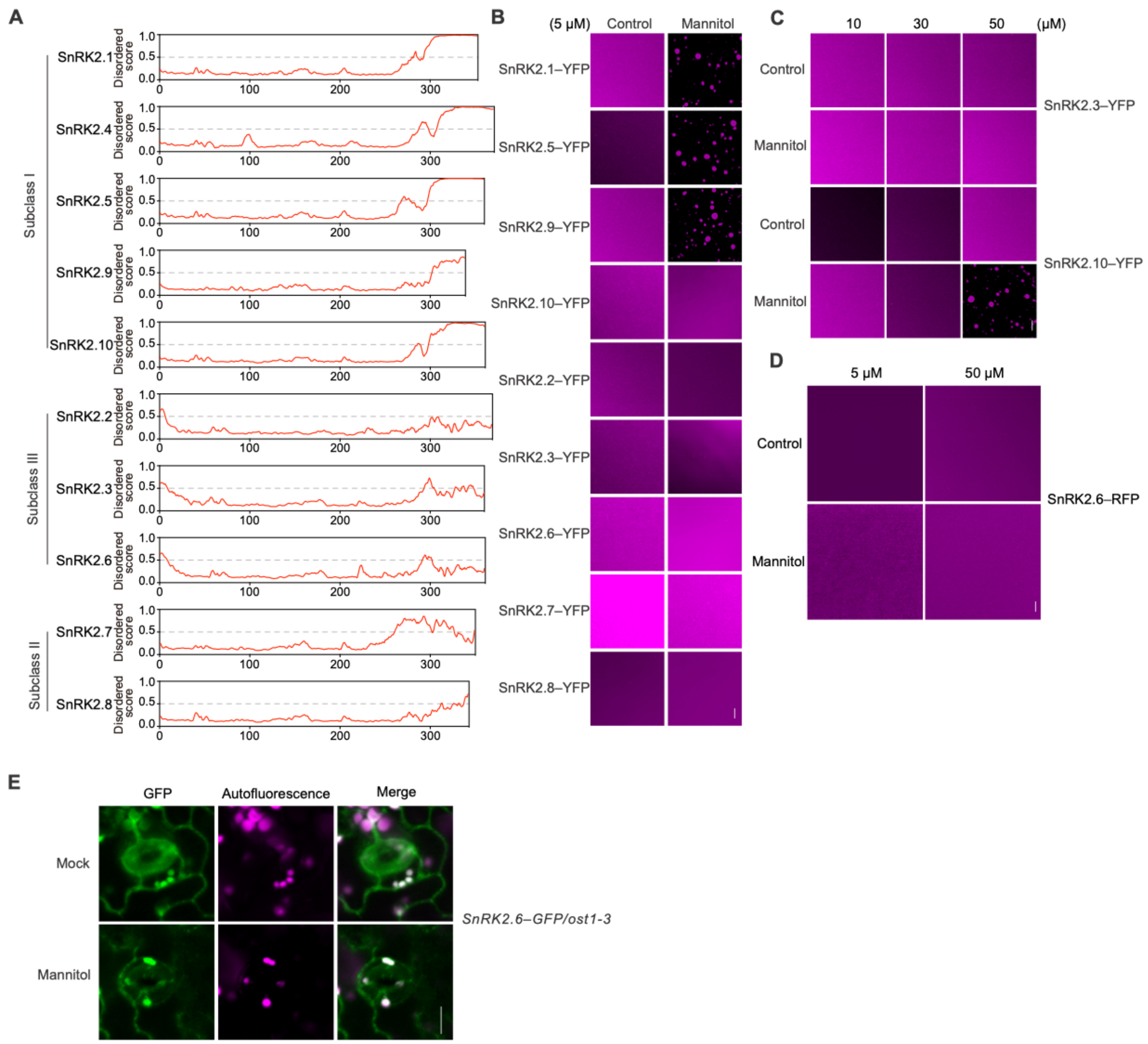


**Figure S6. Predicted IDRs in Arabidopsis SnRK2s and their condensation under osmotic stress. Related to Figure 4.**

(A) The intrinsically disordered regions of SnRK2 proteins were predicted using IUPred2.

(B) *In vitro* condensation assay of 5 µM recombinant SnRK2–YFP proteins.

(C) *In vitro* condensation assay of 10, 30, and 50 µM recombinant SnRK2.3–YFP and SnRK2.10–YFP proteins.

(D) Condensation assay of SnRK2.4–GFP in *SnRK2.4–GFP*/Col-0 Arabidopsis. Images at right are zoomed in from the white boxes at left.

(E) *In vitro* condensation assay of 50 µM recombinant SnRK2.6–YFP protein.

Scale bars: 5 µm in (B)–(D), and 10 µm in (E).

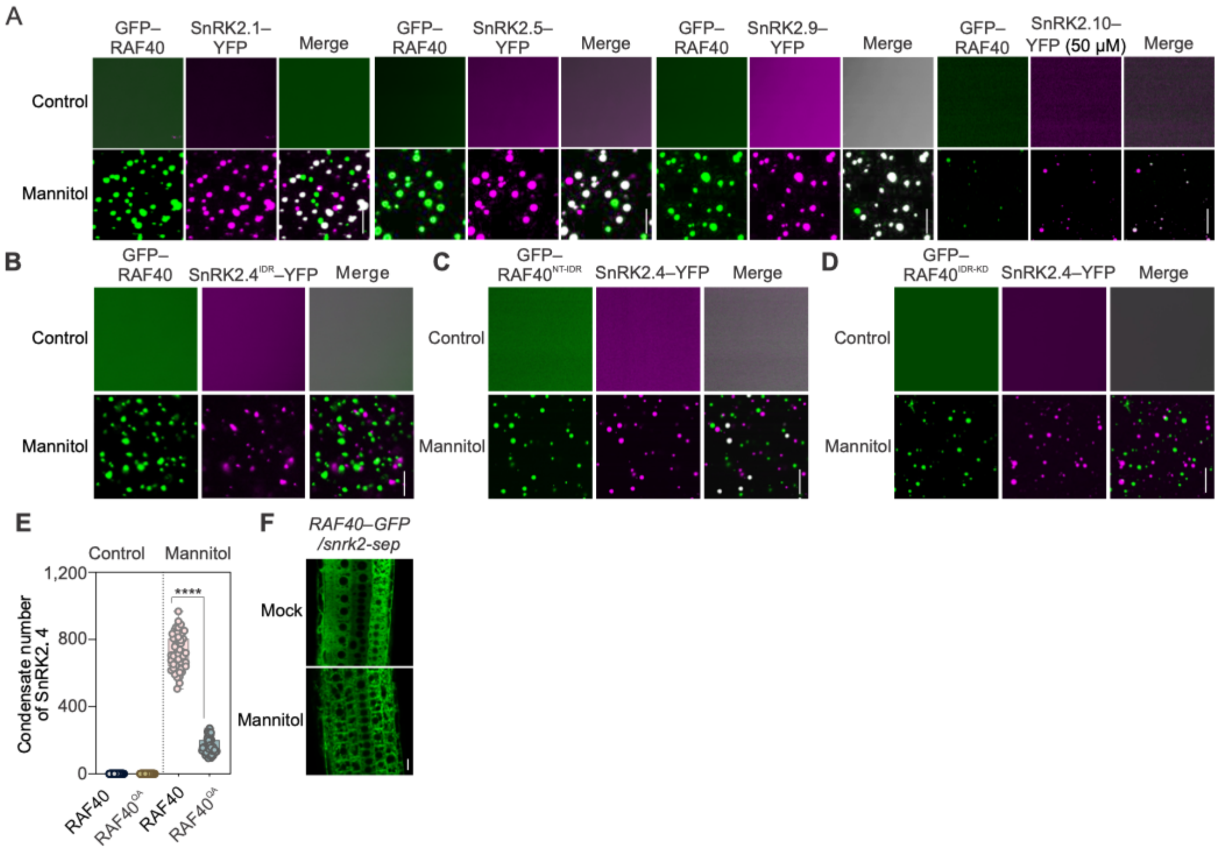


**Figure S7. Co-condensation of GFP–RAF40 and SnRK2–YFP occurs *in vitro* and in *vivo*. Related to Figure 4.**

(A) *In vitro* condensation assay showing the co-condensation of 5 μM GFP–RAF40 with 5 μM SnRK2.1–YFP, 5 μM SnRK2.5–YFP, 5 μM SnRK2.9–YFP, or 50 μM SnRK2.10–YFP, respectively solution without or with 800 mM mannitol.

(B) *In vitro* condensation assay showing the co-condensation of GFP–RAF40 and SnRK2.4$^{IDR}$–YFP in solution without or with 800 mM mannitol.

(C, D) *In vitro* condensation assay showing the co-condensation of SnRK2.4–YFP and GFP–RAF40$^{NT–IDR}$ (C) or GFP–RAF40$^{IDR–KD}$ (D) in solution without or with 800 mM mannitol.

(E) Quantification of the number of SnRK2.4–YFP puncta formed by SnRK2.4–YFP incubated with GFP–RAF40 and GFP–RAF40$^{QA}$ under mannitol treatment. ****, $p < 0.0001$, unpaired two-tailed *t-test* between two groups. Data are means ± SD of n = 9 independent replicates.

(F) Condensation assay of RAF40–GFP in *RAF40–GFP/snrk2-sep* plants in 1/2 MS (Mock) or with 800 mM mannitol. Scale bars: 10 μm. Representative images of n = 3 independent experiments.

(B), (C) and (D) representative images of n = 3 independent experiments. Scale bars: 5 μm in (A)–(D); 10 μm in (F).

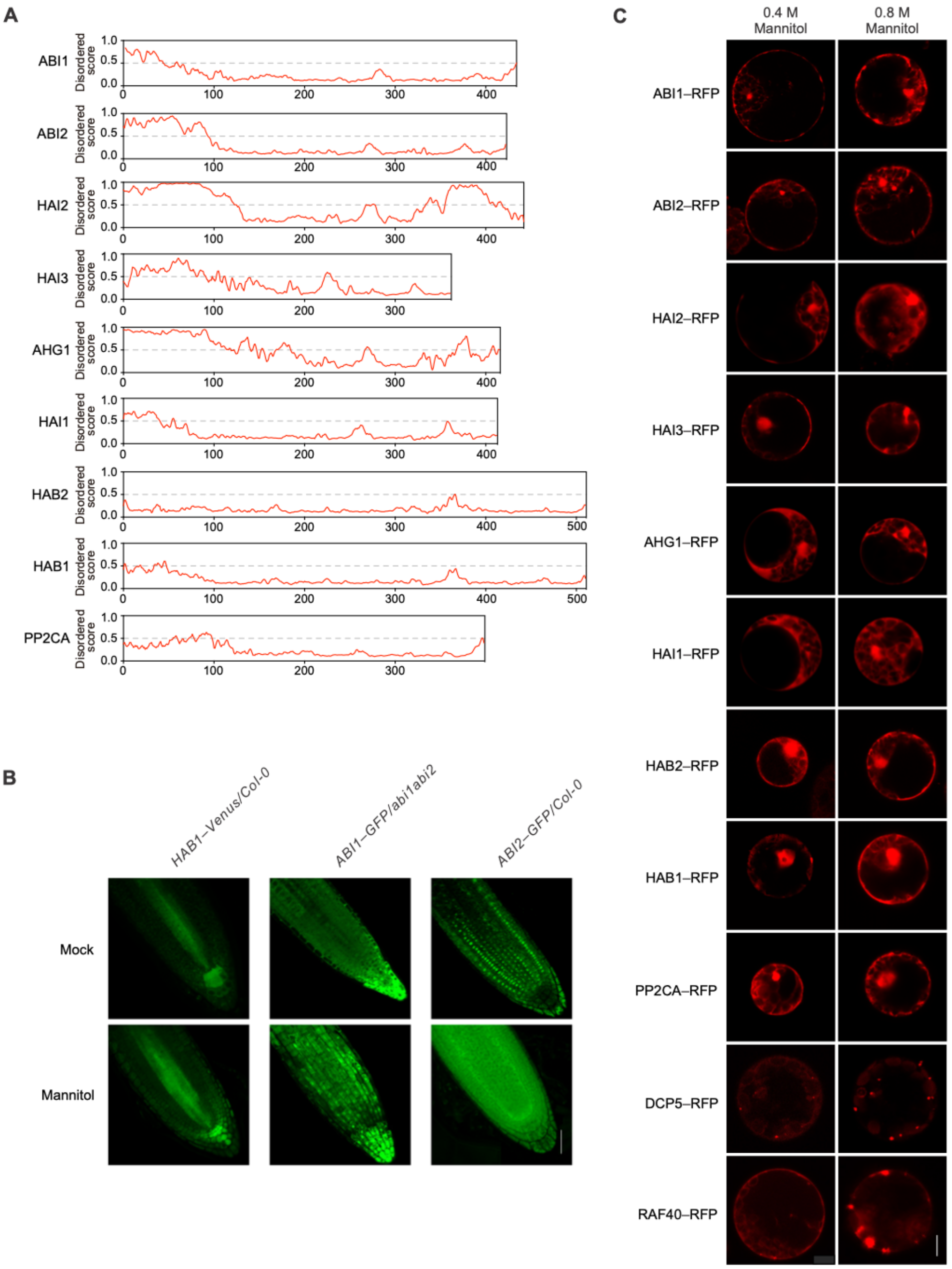


**Figure S8. A-clade PP2Cs do not condense upon mannitol treatment. Related to Figure 5.**
(A) The intrinsically disordered regions of A-clade of PP2Cs were predicted using IUPred2.
(B) Confocal micrographs of transgenic Arabidopsis root cells expressing *HAB1*$_{pro}$*:HAB1–Venus*/Col-0, *ABI1*$_{pro}$*:ABI1–eGFP*/*abi1 abi2*, and *35S*$_{pro}$*:ABI2–GFP*/Col-0 in the presence of 800 mM mannitol.
(C) Confocal micrographs of A-clade PP2Cs transiently expressed in Arabidopsis protoplasts with 400 mM or 800 mM mannitol. DCP5–RFP and RAF40–RFP were used as controls.
Scale bars: 10 µm in (B); 5 µm in (C).

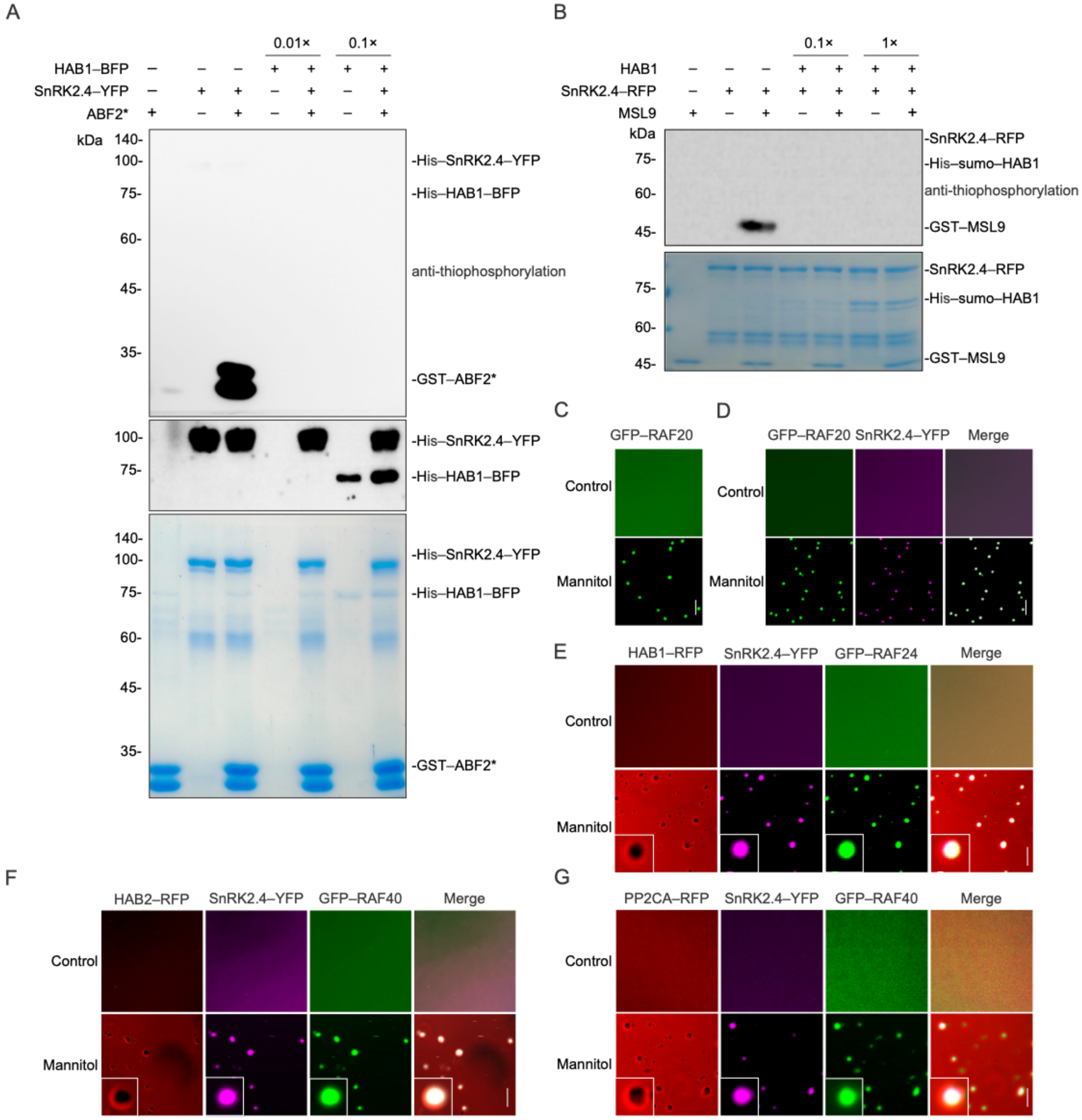


**Fig. S9 A-clade PP2Cs dephosphorylate SnRK2.4 and do not co-condense with RAF24/RAF40 and SnRK2.4 *in vitro*. Related to Figure 5.**

(A) HAB1 inhibits SnRK2.4 activity, in the context of phosphorylation of recombinant GST–ABF2$^*$. Recombinant SnRK2.4 incubated without or with 0.01× or 0.1× HAB1 for 10 min, were used to phosphorylate GST–ABF2* (73–119 aa) in the presence of ATP-γS. Anti γ-S immunoblot (top) indicates the thiophosphorylation of GST–ABF2*, and anti-His and Coomassie staining indicate the abundance of HAB1, SnRK2.4, and GST–ABF2*.

(B) Phosphorylation of an MSL9 fragment[20] by SnRK2.4. SnRK2.4 was dephosphorylated with 0.1× or 1× HAB1 for 10 min, and the MSL9 fragment was used as substrate to indicate the kinase activity of SnRK2.4. Coomassie staining indicates the abundance of HAB1, SnRK2.4, and MSL9.

(C) *In vitro* condensation assay of 5 μM recombinant GFP–RAF20.
(D) *In vitro* condensation assay of 5 μM recombinant SnRK2.4–YFP with 5 μM GFP–RAF20.
(E) *In vitro* condensation assay of 5 μM recombinant SnRK2.4–YFP with 0.5 μM HAB1–RFP and 5 μM GFP–RAF24. Insets for the mannitol treatment are zooms of the indicated small white boxes.
(F) *In vitro* condensation assay of 5 μM recombinant SnRK2.4–YFP with 0.5 μM HAB2–RFP and 5 μM GFP–RAF40. Insets for the mannitol treatment are zooms of the indicated small white boxes.
(G) *In vitro* condensation assay of 5 μM recombinant SnRK2.4–YFP with 0.5 μM PP2CA–RFP and 5 μM GFP–RAF40. Insets for the mannitol treatment are zooms of the indicated small white boxes.
Scale bars: 5 μm in (C)–(G).

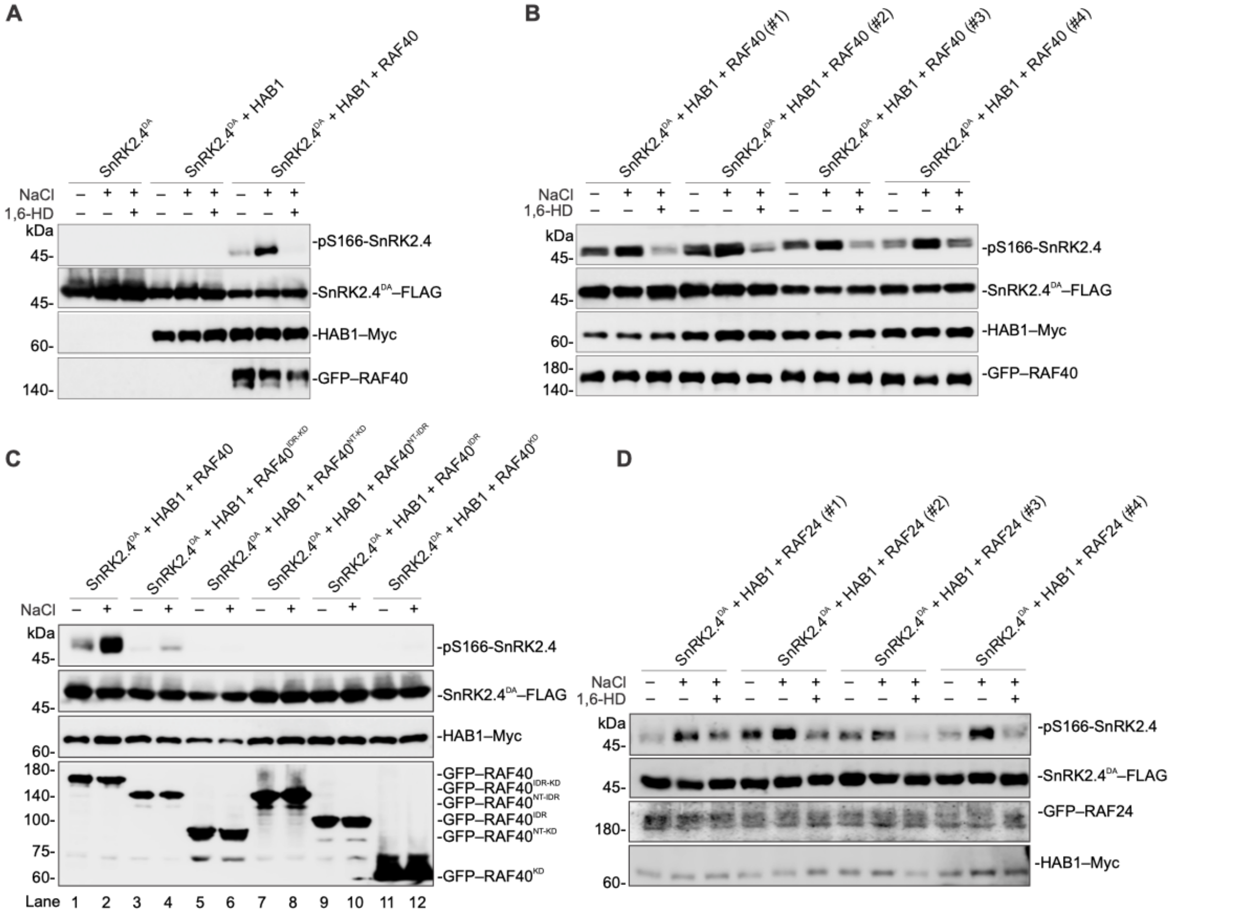


**Figure S10. Reconstitution of the RAF–SnRK2 osmosensing core module in *E. coli*. Related to Figure 6.**

(A) A replicate of the experiment shown in Fig. 6C, immunoblots of pS166-SnRK2.4, SnRK2.4$^{DA}$–FLAG, HAB1-Myc, and GFP–RAF40 abundance in *E. coli* strains with different vector combinations under ± 300 mM NaCl or 5% 1,6-HD for 5 min.

(B) Immunoblots of pS166-SnRK2.4, SnRK2.4$^{DA}$–FLAG, HAB1-Myc, and GFP–RAF40 abundance in four independent clones, in addition to the one shown in Fig. 6C, expressing SnRK2.4$^{DA}$–FLAG, HAB1–Myc, and GFP–RAF40 under ± 300 mM NaCl or 5% 1,6-HD treatment for 5 min.

(C) Immunoblots of pS166-SnRK2.4, SnRK2.4$^{DA}$–FLAG, HAB1–Myc, GFP–RAF40 and its variants in *E. coli* strains expressing SnRK2.4$^{DA}$–FLAG, HAB1–Myc, wild-type RAF40 or RAF40$^{IDR–KD}$, RAF40$^{NT–KD}$, RAF40$^{NT–IDR}$, RAF40$^{IDR}$, RAF40$^{KD}$ with ± 300 mM NaCl treatment for 5 min.

(D) Immunoblots of pS166-SnRK2.4, SnRK2.4$^{DA}$–FLAG, HAB1–Myc, and GFP–RAF24 in four independent *E. coli* clones expressing SnRK2.4$^{DA}$–FLAG, HAB1–Myc, and GFP–RAF24 with ± 300 mM NaCl or 5% 1,6-HD treatment for 5 min.

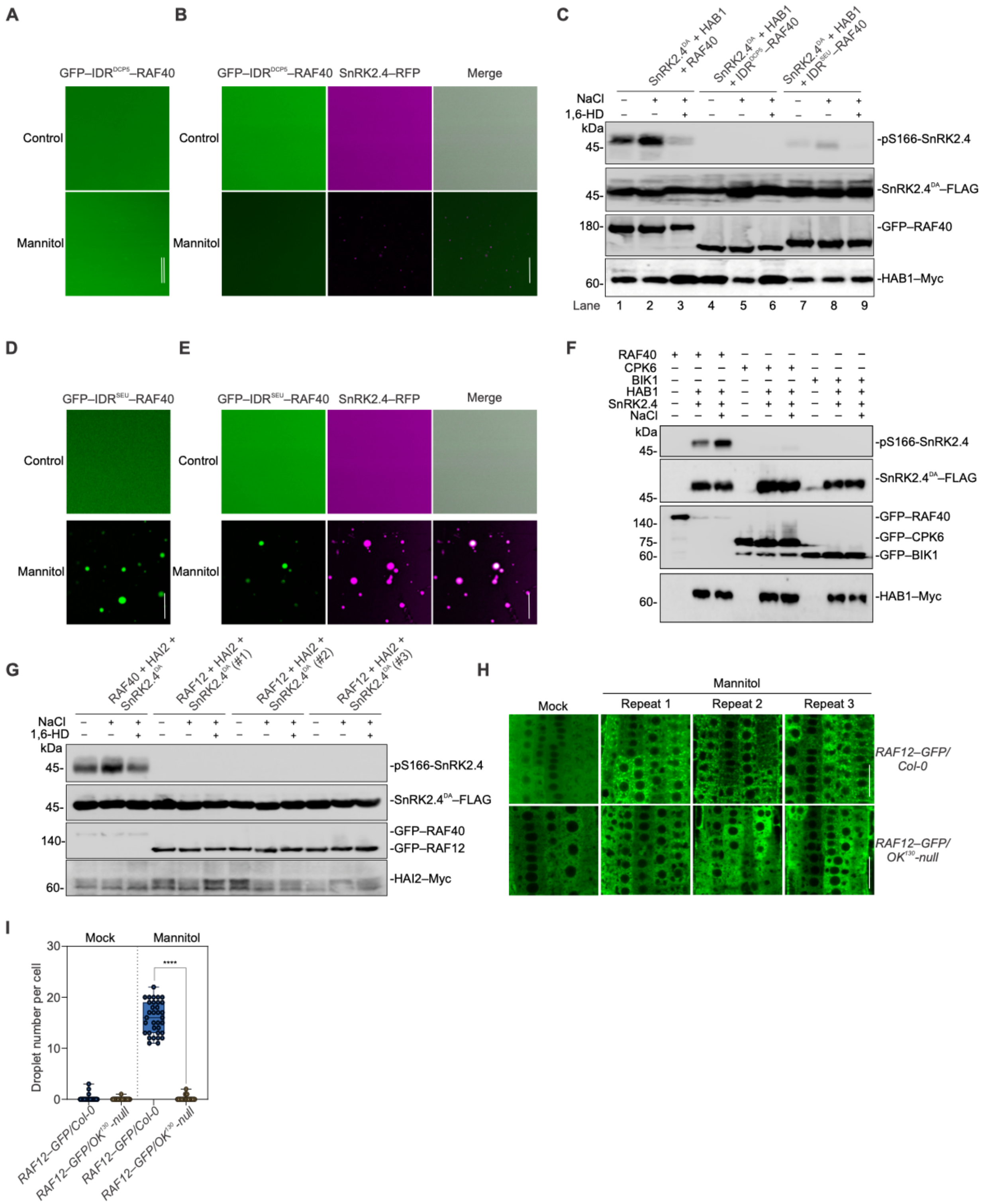


**Figure S11. Condensation and reconstitution assays with RAF40 mutants fused with IDR$^{SEU}$ or IDR$^{DCP5}$, or other kinases reported to activate SnRK2.4. Related to Figure 6.**

(A) *In vitro* condensation assay of 5 μM recombinant GFP–IDR$^{DCP5}$–RAF40.

(B) *In vitro* condensation assay of 5 μM recombinant SnRK2.4–RFP with 5 μM GFP–IDR$^{DCP5}$–

RAF40.
(C) Immunoblots of pS166-SnRK2.4, SnRK2.4$^{DA}$–FLAG, HAB1–Myc, GFP–RAF40/IDR$^{DCP5}$–RAF40/IDR$^{seu}$–RAF40 in *E. coli* strains expressing SnRK2.4$^{DA}$–FLAG, HAB1–Myc, GFP–RAF40 and its variants under ± 300 mM NaCl or 5% 1,6-HD treatment for 5 min.
(D) *In vitro* condensation assay of 5 μM recombinant GFP–IDR$^{SEU}$–RAF40.
(E) *In vitro* condensation of 5 μM recombinant SnRK2.4–RFP with 5 μM GFP–IDR$^{SEU}$–RAF40.
(F) Immunoblots of pS166-SnRK2.4, SnRK2.4–FLAG, HAB1–Myc, and GFP–RAF40/CPK6/BIK1 in *E. coli* strains expressing SnRK2.4$^{DA}$–FLAG, HAB1–Myc, GFP–RAF40/CPK6/BIK1 with ± 300 mM NaCl treatment for 5 min.
(G) Immunoblots of pS166-SnRK2.4, SnRK2.4$^{DA}$–FLAG, HAI2–Myc, GFP–RAF40/ GFP–RAF12 in *E. coli* strains expressing SnRK2.4$^{DA}$–FLAG, HAI2–Myc, GFP–RAF40 or GFP–RAF12 under ± 300 mM NaCl or 5% 1,6-HD treatment for 5 min.
(H) RAF12–GFP condensates in *GFP–RAF12*/Col-0 and *GFP–RAF12/OK$^{130}$-null* plants.
(I) Quantification of condensate numbers per cell in (H). Data are mean ± SD of n = 3 independent replicates. ****, $p < 0.0001$, unpaired two-tailed *t*-test.
Scale bars: 5 μm in (A), (B), (D) and (E); 10 μm in (H).

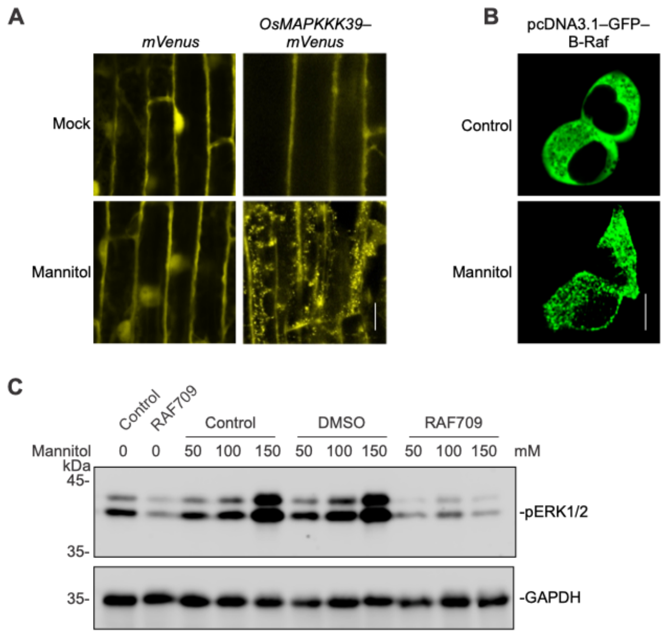


**Figure S12. Mannitol triggers condensation of B4-RAF members in rice and B-Raf in mammalian cells. Related to Figure 6.**
(A) OsMAPKKK39–mVenus fluorescence in $\textit{OsMAPKKK39}_{pro}$*:OsMAPKKK39–mVenus* and mVenus in $\textit{35S}_{pro}$*:mVenus* lines grown in 1/2 MS (Mock) or with 800 mM mannitol.
(B) Confocal micrographs of HEK293T cells transiently expressing pcDNA3.1–GFP–B-Raf after treatment with 100 mM mannitol for 3 min.
(C) Immunoblot showing endogenous ERK1/2 phosphorylation in HEK293T cells with the indicated concentrations of mannitol, without or with the presence of B-Raf inhibitor RAF709. DMSO was used as control for RAF709, and GAPDH was used as the loading control.
Scale bars: 10 µm in (A) and (B).

**Video S1.** Time-lapse imaging showing the recovery after bleaching of RAF40–GFP condensates *in vitro*, related to Figure 1.

**Table S1**. Primers used in this study.
